# Solving High-Dimensional Population Balance Equations via Dynamics-Preserving Autoencoders

**DOI:** 10.64898/2026.08.09.743783

**Authors:** Prateek Gupta, Shourya Verma, Ananth Grama, Doraiswami Ramkrishna

**Affiliations:** Davidson School of Chemical Engineering, Purdue University, West Lafayette, 47907, IN, USA; Department of Computer Science, Purdue University, West Lafayette, 47907, IN, USA

**Author notes:** Corresponding author (D. Ramkrishna). These authors contributed equally to this work (Prateek Gupta, Shourya Verma). (P. Gupta); (S. Verma); (A. Grama).

**Keywords:** Population Balance Equation, Macrophage polarization, Gene Regulatory Networks, Autoencoders, Stochastic Differential Equations, Particle Methods

## Abstract

High-dimensional population balance equations (PBEs) provide a natural framework for modeling heterogeneous cell populations, but their direct numerical solution becomes computationally prohibitive when the internal state space contains many molecular variables. We propose a hybrid mechanistic-machine learning framework for reducing and simulating PBEs defined over high-dimensional intracellular coordinates. The cell population is described by a number density *n*(x, *t*), where x ∈ ℝ^*N*^ represents gene and protein states associated with macrophage activation. A dynamics-preserving autoencoder maps this state space to a low-dimensional latent coordinate **z** ∈ ℝ^*d*^, with *d* ≪ *N*, while retaining key qualitative features of the underlying gene regulatory network, including attractor structure and multistability. Mechanistic information from the original regulatory dynamics is used to construct interpretable drift and diffusion terms for the reduced latent-space PBE. The reduced PBE is solved using a stochastic Lagrangian particle representation, in which particles evolve according to stochastic differential equations (SDEs) corresponding to the latent drift and diffusion fields. The resulting latent-space solution is subsequently decoded and propagated back into the original state space to recover physically interpretable cellular dynamics. We demonstrate the framework on macrophage polarization under cytokine-dependent regulation, including gene knockout perturbations. Overall, the proposed framework provides a computationally tractable and mechanistically interpretable route for integrating single-cell genomic data with population balance models of cell-state dynamics.

## 1. Introduction

In this article, we investigate a biological problem that is not a customary component of chemical engineering. It, however, earns a rightful place in this literature not only because of its strong link to the analysis of chemical reactions, but also because of the availability of chemical engineering tools that can profoundly address issues crucial to the solution of the problem. The problem of interest is the phenomenon of macrophage polarization. Past efforts on modeling have sought the aid of kinetic models, based on average cell behavior without regard to the simultaneous presence of numerous macrophages ranging between phenotype *M*_1_ (pro-inflammatory, host-defense) and *M*_2_ (anti-inflammatory, tissue repair).

Tissue homeostasis emerges as a consequence of multiple cell populations working in harmony to maintain a stable internal environment (Parigini and Greulich, 2024; Frankel and Lim, 2018). Cells differentiate, proliferate, die, or polarize among other fates to keep a strict balance of their numbers. This balance is intriguingly delicate and robust at the same time - self-correcting mechanisms endow the tissue with resilience to normal environmental variations, yet, may leave it vulnerable to specific, extreme disruptions (Meizlish, Franklin, Zhou and Medzhitov, 2021; Pardee, 2006). Wounds, in particular, are a specific example of disruption where the skin barrier is compromised, leaving the underlying tissue both damaged and prone to pathogenic insult (Jiao, Zhi, You, Wang, Wu and Jia, 2024; Rosińczuk, Taradaj, Dymarek and Sopel, 2018). Besides a few approved treatments, most other wound therapies are in various stages of the pipeline, and the most advanced treatments aim to manipulate cell types (Mao, Chen, Cai, Qian, Liu, Zhao, Zhang, Sun and Cui, 2022; Knoedler, Knoedler, Kauke-Navarro, Rinkevich, Hundeshagen, Harhaus, Kneser, Pomahac, Orgill and Panayi, 2023; Huelsboemer, Knoedler, Kochen, Yu, Hosseini, Hollmann, Choi, Stögner, Knoedler, Hsia et al., 2024).

Among the top contenders for phenotype manipulation are macrophage cells, which are specialized immune cells with very high degree of plasticity and extreme diversity (Chen, Liu, Tang, Luo, Liang and He, 2023a; Chen, Wu, Wang, Zhang, Song, Fu, Kong and Shi, 2023b; Sica, Mantovani et al., 2012). Equipped with super-sensing capabilities, they can rapidly sample their milieu and polarize into functionally distinct subsets (Qian, Yun, Yao, Cao, Liu, Hu, Zhang and Luo, 2019). While originally viewed as cells with binary fate commitment (M1 or M2), recent advances in single-cell omics have updated their phenotypic status to a spectrum (Mosser and Edwards, 2008; Murray, 2017). Exhibiting a range of activation states, macrophages can lean toward a pro-inflammatory (M1-like) or anti-inflammatory (M2-like) profile, or occupy any intermediate position along this functional spectrum (Wang, Liang and Zen, 2014; Muñoz-Rojas, Kelsey, Pappalardo, Chen and Miller-Jensen, 2021). In rare instances, they can even defy this spectral categorization (Szulzewsky, Pelz, Feng, Synowitz, Markovic, Langmann, Holtman, Wang, Eggen, Boddeke et al., 2015; Kalish, Lyamina, Manukhina, Malyshev, Raetskaya and Malyshev, 2017).

Ordinary differential equations (ODEs) are commonly used to model the aggregate behavior of cell populations, owing to their computational efficiency and ability to capture continuous-time dynamics (Alon, 2023). Agent-based models (ABM) are another popular choice for modeling small-sized populations using boolean logic (Minucci, Heise and Reynolds, 2024; Nickaeen, Ghaisari, Heiner, Moein and Gheisari, 2019). Additionally, bioinformatic approaches can process vast amounts of data to identify complex correlations within cell populations, but often offer no mechanistic insight (Li, Menoret, Farragher, Ouyang, Bonin, Holvoet, Vella and Zhou, 2019; Henlon, Panir, McIntyre, Hogg, Dhami, Cuff, Senior, Moolchandani-Adwani, Courtois, Horne et al., 2024). Recent Machine Learning (ML) frameworks have leveraged data to make cell-fate predictions at population level (Fischer, Fiedler, Kernfeld, Genga, Bastidas-Ponce, Bakhti, Lickert, Hasenauer, Maehr and Theis, 2019; Zheng, Barile, Wilson, Huang, Theis and Göttgens, 2025; Lange, Bergen, Klein, Setty, Reuter, Bakhti, Lickert, Ansari, Schniering, Schiller et al., 2022; Weiler, Lange, Klein, Pe’er and Theis, 2024). They solve an inverse problem of identifying the (time- and state-dependent) coefficients of a parabolic PDE that tracks cell number densities *n*(x, *t*). However, in the absence of time-resolved datasets, the inference of these coefficients become non-unique and often ill-posed (Weinreb, Wolock, Tusi, Socolovsky and Klein, 2018). Furthermore, the coefficients are frequently modeled as splines (or using Radial Basis Functions, RBF) and lack any kinetic or biochemical interpretation. Many of these studies further disregard the important dynamical features such as multi-stability and bifurcations.

In this paper, we propose a new method that not only overcomes the aforementioned limitations, but also opens up new avenues for *in silico* drug testing and therapeutic developments. The biological system of interest is a population of heterogeneous macrophages whose phenotypic evolution is studied through the lens of a hybrid Population Balance Model/ Framework (hPBM/hPBF). PBMs have been used extensively in the domain of chemical engineering to model particulate systems (Ramkrishna, 2000; Ramkrishna and Singh, 2014; Ramkrishna and Mahoney, 2002; Bhole, Joshi and Ramkrishna, 2008; Ramkrishna, 2005) and their strengths are leveraged here to show how a highly heterogeneous set of macrophages evolve under the dynamic influence of gene-environment (*G* × *E*) interactions. We use the manifold hypothesis (Fefferman, Mitter and Narayanan, 2016) and the model-reduction capabilities of autoencoders (Berahmand, Daneshfar, Salehi, Li and Xu, 2024) to project a high-dimensional PBE onto a low-dimensional latent space, where the governing dynamics are solved using a stochastic particle algorithm coupled to the autoencoder. Our architecture further integrates the SINDy framework (Brunton, Proctor and Kutz, 2016a) and introduces novel loss functions tailored to preserve the underlying dynamics. To the best of our knowledge, this is among the first studies to successfully handle PBEs in very high dimensions (up to 24 dimensions) with scalability beyond.

## 2. Background and Methodology

### 2.1. System Description

Macrophages sense and integrate complex microenvironmental stimuli to orchestrate transitions across their phenotypic landscape. We refer to these transitions as ‘polarization’. The underlying circuitry (Fig. 1A) enabling these transitions is called a Gene Regulatory Network (GRN), which comprises mainly of regulators (e.g., transcription factors (TFs)) and associated targets. As discussed in §1, phenotypes M1 and M2 represent the specialized extremes of the polarization spectrum. The GRN governing these programs exist in a state of dynamic antagonism, characterized by complex positive and negative feedback loops that preclude the existence of ‘pure’ M1 or M2 states. The traditionally defined subsets (e.g., M2a, M2b, M2c) emerge as distinct regions within this high-dimensional landscape, representing varying degrees of transcriptional overlap rather than discrete, mutually exclusive categories. A core subset of 12 genes - STAT1, STAT6, IRF5, HIF1*α*, NF*κ*B, JMJD3, IRF4, KLF4, PPAR*γ*, STAT3, IL-1*β*, and IL-10 – implicated in the polarization process is utilized to construct a kinetic model of gene expression (Marku, Verstraete, Raynal, Madrid-Mencía, Domagala, Fournié, Ysebaert, Poupot and Pancaldi, 2020). Each gene is represented by its normalized mRNA concentration *m*_*i*_ ∈ [0, 1] and protein concentration *p*_*i*_ ∈ [0, 1], yielding a 24-dimensional internal state vector x = [**m**^⊤^, **p**^⊤^]^⊤^ ∈ [0, 1]^24^. The normalization bounds are *m*_max,*i*_ = (*α*_*i*_ + *γ*_*i*_)*/δ*_*i*_ and *P*_max,*i*_ = *β*_*i*_*m*_max,*i*_*/µ*_*i*_, giving dimensionless ODEs

**Figure 1.**
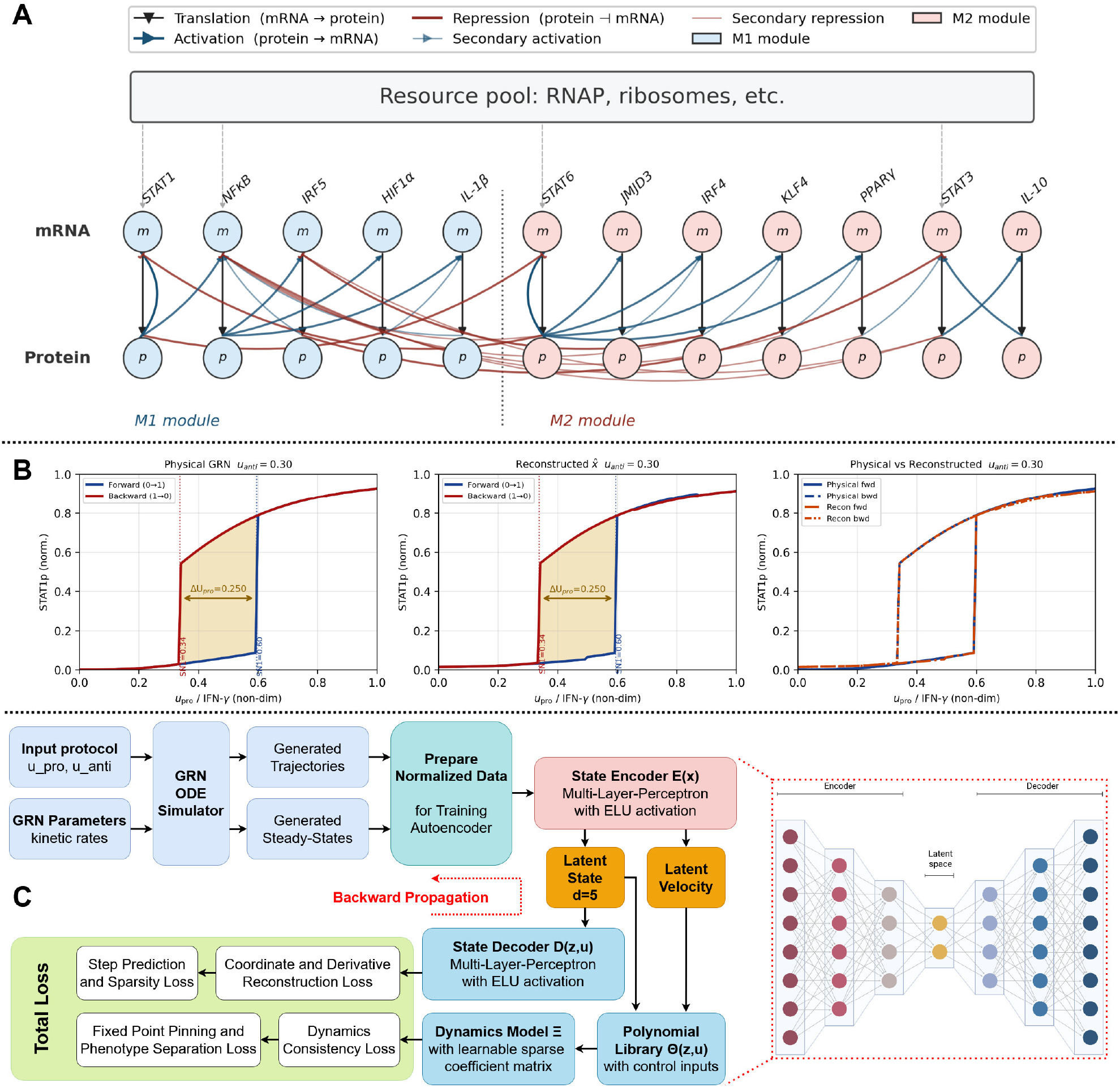
Gene regulatory network model and autoencoder training pipeline for macrophage polarization. **(A)** Schematic of the 12-gene GRN governing macrophage M1/M2 polarization (containing 12 Genes and 12 Proteins). Genes are partitioned into an M1 module (STAT1, NF*κ*B, IRF5, HIF1*α*, IL-1*β*; blue) and an M2 module (STAT6, JMJD3, IRF4, KLF4, PPAR*γ*, STAT3, IL-10; salmon) coupled by a mutual repression toggle. Bold blue and red arrows denote activation and repression; lighter arrows indicate secondary interactions. **(B)** At fixed (*u*_anti_ = 0.30), the full GRN hysteresis loop in STAT1p is reproduced by continuing the learned latent SINDy dynamics and decoding the resulting branches to reconstructed physical space, 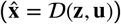. Agreement in the forward/backward switching thresholds and hysteresis width (Δ*U*_pro_) indicates that the reduced model preserves the bistable switching structure of the original GRN. **(C)** Autoencoder pipeline. The GRN ODE simulator generates trajectory and steady-state data under varied input protocols, normalized and batched for training. The encoder *ε*(x) maps the 24-dimensional state to a *d*-dimensional latent state **z** and, via a chain-rule Jacobian-vector product, to the latent velocity ż. A polynomial library **Θ**(**z, u**) combined with a learnable sparse coefficient matrix **Ξ** predicts latent dynamics as ż = **Θ**(**z, u**)**Ξ.** The decoder *D*(**z, u**) reconstructs the full state and its derivative via a second Jacobian-vector product. The total loss combines reconstruction, dynamics consistency, one-step prediction, sparsity, fixed-point pinning, and phenotype separation terms, with gradients backpropagating jointly through all components (dashed red arrow). Inset: encoder-latent-decoder MLP architecture.

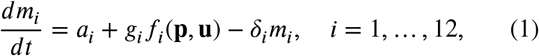

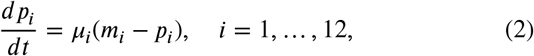

where *a*_*i*_ = *α*_*i*_*/m*_max,*i*_ and *g*_*i*_ = *γ*_*i*_*/m*_max,*i*_ are normalized basal and inducible production rates (Gupta and Ramkrishna, 2026), and the protein ODE simplifies exactly because *β*_*i*_*m*_max,*i*_*/P*_max,*i*_ = *µ*_*i*_ by construction. The transcriptional regulation functions *f*_*i*_ are products of Hill activation and repression terms,

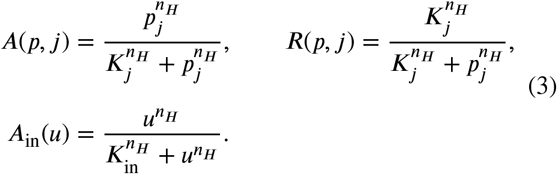

with Hill coefficient *n*_*H*_ = 3 and input threshold *K*_in_ = 0.5. Physical parameters *δ*_*i*_, *µ*_*i*_, *β*_*i*_ are taken from TTDB (Jiang, Xu, Li, Wang and Xie, 2026) and CHX-chase measurements, and the normalized Hill threshold *K*_nd,*i*_ = *K*_*p*_*/P*_max,*i*_ is computed from *K*_*p*_ = 1200 (physical protein units). Full model specification is given in Supplementary file S1.

### 2.2. Population Balance for Cell Populations

For a cell population in which each cell is characterized by an internal state vector x ∈ Ω ⊂ ℝ^*N*^, the number density *n*(x, *t*) satisfies the population balance equation

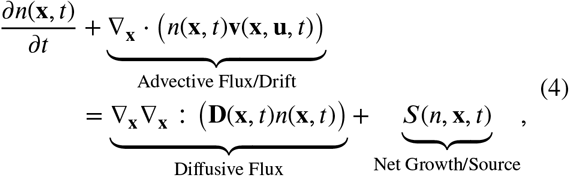

where **v :** Ω → ℝ^*N*^ is the deterministic drift given by the intracellular GRN ODEs ((1) & (2)) and **D** ∈ ℝ^*N*×*N*^ is the anisotropic diffusion tensor encoding molecular noise.

For *N* = 24, both the grid-based discretization and direct Monte Carlo simulation of Eq. (4) at biologically meaningful ensemble sizes are computationally prohibitive. Particle methods address challenges associated with high-dimensional grids, but incur the cost of integrating the full *N*-dimensional ODE per particle per step (Chertock, 2017). Macroscopic population observables, including phenotype fractions and marker distributions measured by flow cytometry, correspond to moments or marginals of *n*(x, *t*). In general, the *k*^th^ moment of the population density is defined as:

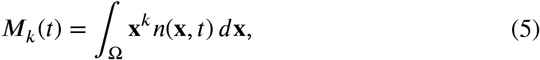

with lower-order moments encoding quantities such as the total population size, mean phenotype, and population variability. Computing these quantities requires solving Eq. (4), which motivates the dimension reduction strategy developed in this work.

### 2.3. Kinetically Constrained Diffusion

The chemical Langevin equation (CLE) provides a diffusion approximation to the discrete stochastic chemical kinetics of a reaction network when molecular copy numbers are large enough for a continuous approximation but small enough that fluctuations are non-negligible (Gillespie, 2000). For the normalized GRN state x = [**m**^⊤^, **p**^⊤^]^⊤^ ∈ [0, 1]^24^ consisting of normalized mRNA **m** ∈ [0, 1]^12^ and protein **p** ∈ [0, 1]^12^, the diffusion tensor **D**(x, *t*) is diagonal with entries

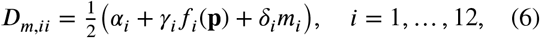

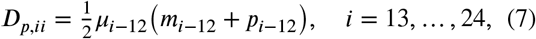

where *α*_*i*_, *γ*_*i*_, *δ*_*i*_, *µ*_*i*_ are normalized production, inducible production, degradation, and translation rates, and *f*_*i*_(**p**) is the transcriptional regulation function for gene *i* (defined in Section 2.1). The diagonal structure reflects the independence of the Poisson noise sources for each transcription and translation reaction and off-diagonal terms vanish under the diffusion approximation. This diagonal CLE covariance is computed analytically from GRN parameters and the current state, requiring no additional fitting. (See supplementary file S1 for full derivation of the tensor).

### 2.4. Model Order Reduction via Deep Learning

Solving a 24-dimensional PBE directly is computationally intractable and motivates the use of model order reduction techniques. This is further supported by the manifold hypothesis, which suggests that high-dimensional data generated by complex systems often concentrate on or near a lower-dimensional subspace. In the present setting, this view is consistent with the observation that GRN dynamics often evolve on low-dimensional attractors associated with stable cell fates (Zhu, Zhang, Sun, Dai, Wen, Zhou and Chen, 2026). In the context of macrophage polarization, these attractors correspond to pro-inflammatory (M1-like) and anti-inflammatory (M2-like) regimes, suggesting an underlying low-dimensional structure governing phenotype transitions. As a preliminary qualitative check, one may examine linear projections such as PCA on trajectories generated from the GRN ODE model under multiple initial conditions. However, since PCA is a linear method, it cannot fully resolve nonlinear manifold structure, curved manifolds, or basin geometry, and thus serves only as a limited visual indicator rather than the basis of the reduction. This motivates the task of learning the underlying low-dimensional manifold directly using deep learning methods (Chui and Mhaskar, 2018; Whiteley, Gray and Rubin-Delanchy, 2026).

#### 2.4.1. Autoencoder Model

Learning a low-dimensional manifold is equivalent, in our setting, to learning a closed set of governing ODEs in the latent *z*-space. Sparse Identification of Nonlinear Dynamics (SINDy) provides a deep learning framework for this task by identifying sparse representations of dynamical systems (Brunton et al., 2016a; Kaheman, Kutz and Brunton, 2020). It identifies governing equations of the form ż = **f** (x) by selecting a sparse subset of candidate functions from a library Θ(x) via sequentially thresholded least squares (or STLSQ):

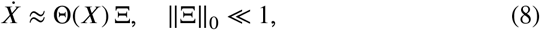

where 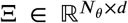 is the sparse coefficient matrix whose columns specify, for each latent coordinate, which library terms contribute to its time derivative and with what weight. SINDy assumes sparsity in the measurement coordinates. To address cases where sparsity holds only in a transformed space, a SINDy autoencoder was proposed (Champion, Lusch, Kutz and Brunton, 2019) that jointly learns an encoder *ε* : ℝ^*N*^ → ℝ^*d*^ and decoder *D* : ℝ^*d*^ → ℝ^*N*^, together with sparse latent dynamics. The latent variables **z** = *ε* (x) are constrained to satisfy 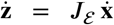, where *J*_ε_ = *∂ε/∂*x ∈ ℝ^*d*×*N*^ is the Jacobian of the encoder, the matrix of partial derivatives that linearly maps an observed state-space velocity 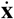 into the corresponding latent-space velocity via the chain rule 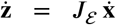. This framework simultaneously discovers a nonlinear coordinate system and sparse governing equations in latent space. Extensions to include control inputs (SINDy-c) augment the library with forcing terms *u*_*j*_ and bilinear terms *z*_*i*_*u*_*j*_ (Brunton, Proctor and Kutz, 2016b).

In this work, we adopt this extension and additionally condition the decoder on the control input **u**. Our autoencoder maps the normalized GRN state x ∈ [0, 1]^24^ through three networks: an encoder ε : ℝ^24^ → ℝ^*d*^, a dynamics model *f*_Ξ_ : ℝ^*d*^ × ℝ^2^ → ℝ^*d*^, and an input-conditioned decoder *D* : ℝ^*d*^ × ℝ^2^ → ℝ^24^. Both *ε* and *D* are fully connected networks with ELU activations and two hidden layers of width 128. The decoder takes [**z**^⊤^, **u**^⊤^]^⊤^ as input, making the reconstruction 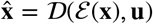 explicitly dependent on the cytokine input. This is essential because the same latent coordinate encodes distinct physical gene expression states under different signaling environments. We train with latent dimension *d* ∈ {2, 3, 4, 5, 6} and report results for *d* = 5, which achieves reconstruction error below 10^−3^ while retaining the sparsity needed for interpretable dynamics.

#### 2.4.2. Polynomial Library with Control Inputs

The dynamics model evaluates a library of candidate terms 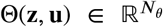 and computes the predicted latent velocity as:

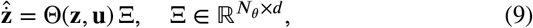

where Ξ is a learnable sparse coefficient matrix with a binary mask 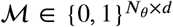 that is progressively pruned during training. The library contains: (i) a constant term; (ii) all monomials 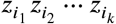 of degree *k* ≤ *q* in the *d* latent coordinates, enumerated via combinations with replacement; (iii) the two control inputs *u*_pro_, *u*_anti_; and (iv) all bilinear cross terms *z*_*i*_*u*_*j*_. For *d* = 5 and polynomial order *q* = 3, the library has 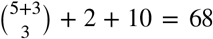 features per equation, for a total of 68 × 5 = 340 learnable coefficients before masking. This library is strictly more expressive than the original SINDy-c library in the latent coordinates, and the sparsity promotion during training recovers the active terms. (See supplementary file S1 for expanded library).

#### 2.4.3. Training Losses

Training is the process of adjusting the free numerical parameters of the encoder, decoder, and dynamics model so that their collective output best matches observed data. A scalar-valued loss function measures the total mismatch between model predictions and data. An optimization algorithm then iteratively updates all parameters in the direction that reduces this loss, repeating over many passes through the dataset until the loss converges. Training minimizes a composite loss over batches of trajectory snapshots 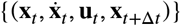:

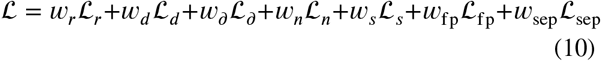

The first five terms follow Champion et al. (2019). The reconstruction loss *L*_r_ = ‖ *D* (*ε*(x) **u**) − x‖^2^ penalizes coordinate-space reconstruction error. The dynamics consistency loss

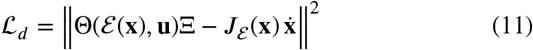

enforces that the autoencoder velocity matches the true latent velocity obtained by pushing 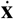 through the encoder Jacobian *J*_*ε*_ = *∂ε/∂*x, computed via forward-mode automatic differentiation (JVP). The derivative reconstruction loss 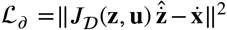 enforces that decoding the predicted latent velocity recovers the original state-space velocity. The one-step prediction loss 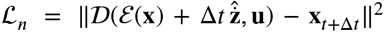 provides step consistency. The sparsity loss ℒ = ‖ Ξ ⊙ ℳ ‖_1_ promotes sparse equations.

The two novel losses extend the framework to mechanistically constrained coordinate learning. The *fixed-point pinning loss*

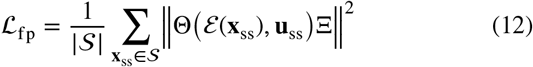

forces the autoencoder velocity field to vanish at the encoded positions of the GRN’s stable steady states *S*, which are precomputed by integrating the deterministic GRN ODE to convergence from a Latin hypercube of initial conditions. This anchors the learned dynamics to the known attractor structure of the mechanistic model. The *phenotype separation loss*

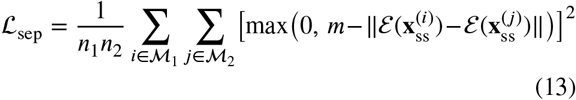

is a hinge margin loss over all M1-M2 pairs of encoded steady states within a batch, with margin *m*. It forces the encoder to place the two attractors in geometrically distinct regions of latent space, ensuring that the learned coordinate system reflects the multistable topology of the GRN. Standard autoencoders prioritize reconstruction efficiency, frequently distorting the underlying dynamical topology (Bank, Koenigstein and Giryes, 2023). Our novel fixed-point pinning and phenotype separation losses explicitly counteract this by forcing the neural network to preserve the mechanistic attractor basins, bifurcations, and geometric boundaries of the original GRN. This ensures that the distinct M1 and M2 biological fates remain topologically separated in the latent space.

Training proceeds in two phases: a reconstruction warmup of 100 epochs with only ℒ_*r*_ active, allowing the encoder and decoder to establish a faithful coordinate map before the dynamics losses are introduced. This is followed by 400 epochs of joint training with all seven losses and a ReduceLROnPlateau scheduler. Thresholding of Ξ at value λ = 10^−2^ is applied every 50 epochs during joint training to enforce sparsity.

### 2.5. PBE Transformation under the Autoencoder

The autoencoder architecture described in §2.4.1 defines an encoding map *ε* : *X* → *Z*, where *X* ⊂ ℝ^24^ denotes the original GRN state space and *Z* ⊂ ℝ^*d*^, with *d* ≪ 24, denotes the learned latent space. For a given input condition **u**, the latent coordinates are defined by **z** = *ε*(x). The decoder map is denoted by 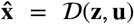. We now show how the PBE transforms under this nonlinear map. For each latent coordinate *z*_*i*_ = *ε*_*i*_(x), we define the encoder Jacobian^1^ with respect to the state^*i*^ variables as:

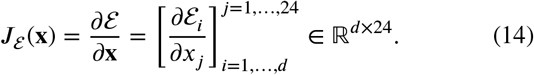

And the Hessian of the encoder as

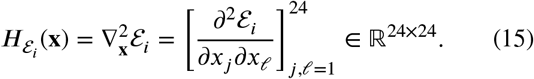

If the high-dimensional parabolic operator in (4) is transformed into latent coordinates, the **latent number density** ñ (**z**, *t*) satisfies the reduced drift-diffusion PBE:

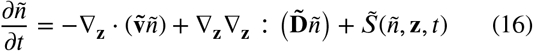

where the tilde sign denotes a function is in the latent space. The transformed diffusion tensor is given by local covariance propagation through the encoder as:

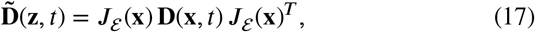

and the transformed drift contains two contributions:

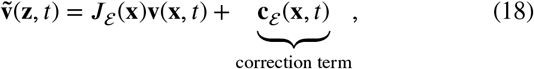

where the *i*-th component of the correction term is given by:

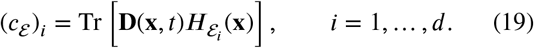

This correction arises because a nonlinear coordinate transformation of a second-order parabolic operator produces an additional drift-like term involving the curvature of the transformation (see Supplementary S1). The source or growth term is transformed by evaluating it on the decoded state: 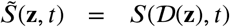. In this work, the SINDy-autoencoder is trained on deterministic GRN trajectories and so the learned latent drift satisfies:

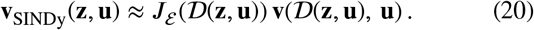

Therefore, if the reduced PBE is intended to approximate the transformed high-dimensional parabolic operator, the drift used in the latent PBE should be written as

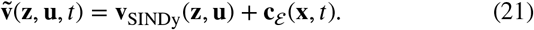

Alternatively, if the latent diffusion is treated as an effective reduced-space closure rather than an exact pushforward of the original diffusion operator, the correction term may be neglected initially and assessed later through sensitivity analysis.

### 2.6. Solving Latent-space PBE

Equation (16) is to be solved numerically within the latent space of the autoencoder (AE). We note that the conservative drift-diffusion part of (16) admits the following microscopic stochastic representation (Gardiner, 2009):

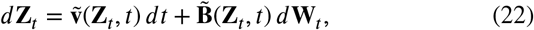

where **Z**_*t*_ ∈ ℝ^*d*^ is a stochastic particle trajectory in the *d*- dimensional latent space, *d***W**_*t*_ is a standard vector Wiener process, and the noise-amplitude matrix 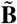 satisfies

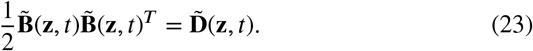

Thus, rather than discretizing the latent number density (ñ) on a mesh, we can approximate it by an ensemble of particles (Chertock, 2017),

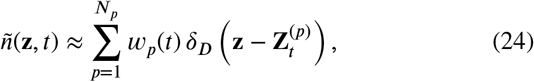

where *δ*_*D*_(.) is the Dirac delta distribution, 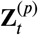 is the position of particle *p, w*_*p*_ (*t*) is its weight, and *N*_*p*_ is t^*t*^he total number of particles. The weights are updated ac^*p*^cording to:

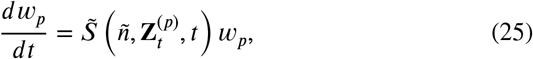

and the particle trajectories are advanced in the latent space using the Euler-Maruyama discretization of (22) as:

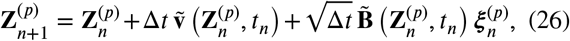

where 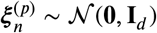 is a standard Gaussian random vector in the latent space (Higham, 2001; Kloeden and Platen, 1992). An important distinction between the physical-space PBE and the latent-space PBE is that the latent-space transport coefficients are not available in closed form. In the original physical state space, drift and diffusion are prescribed by the mechanistic gene-regulatory model (eqns. (1), (2) & (6)). However, after transformation through the nonlinear encoder **z** = *ε* (x), the corresponding latent-space PBE coefficients depend on derivatives of the trained autoencoder.

## 3. Results

We now investigate the fidelity of the coupled autoencoder-particle solver framework. The main objective is to determine whether the latent-space formulation remains consistent with our intuition about macrophage phenotypic dynamics under different perturbations. For instance, if macrophages are stimulated exclusively with M1-associated cytokines, then the population should progressively evolve toward the M1 phenotypic state. Reproducing such behavior is important because it indicates that the coupled framework preserves the expected biological response of the original GRN. Since the framework is ultimately intended to study the dynamical systems behavior of macrophage gene regulatory networks at the population level, we additionally investigate whether key dynamical features, including multistability, bifurcations, and hysteresis, are retained within the latent representation or not. All simulations in this section correspond to a closed in vitro population with 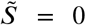. Thus, the reported population dynamics reflect redistribution of macrophages across phenotypic state space rather than changes in total cell number due to recruitment, proliferation, or clearance.

### 3.1. Ablation Study and Dimensionality Selection

The dimensionality sweep in Fig. 2A highlights the importance of choosing a latent dimension that is large enough to represent the attractor geometry but small enough to permit sparse and stable dynamics identification. At *d* = 2, the surrogate achieves only 46% region agreement (40/87) and nearly eliminates the bistable regime, indicating that two latent coordinates are insufficient to represent the M1/M2 toggle together with the downstream regulatory structure. Increasing the dimension to *d* = 3 improves agreement to 74% (64/87), but a substantial portion of the bistable interior remains misclassified, suggesting that the attractor locations can be partially represented while their basin structure remains unresolved. The *d* = 4 and *d* = 6 models perform worse, with 52% (45/87) and 23% (20/87) agreement, respectively. These failures suggest that increasing latent dimension does not monotonically improve dynamical fidelity: larger latent spaces expand the SINDy library and may admit spurious polynomial dynamics that fit local trajectory velocities but fail to preserve the global fixed-point structure. In contrast, *d* = 5 provides the best balance between geometric expressivity and sparse dynamics identification, recovering monostable M1, monostable M2, and bistable regimes with 93.1% agreement.

**Figure 2.**
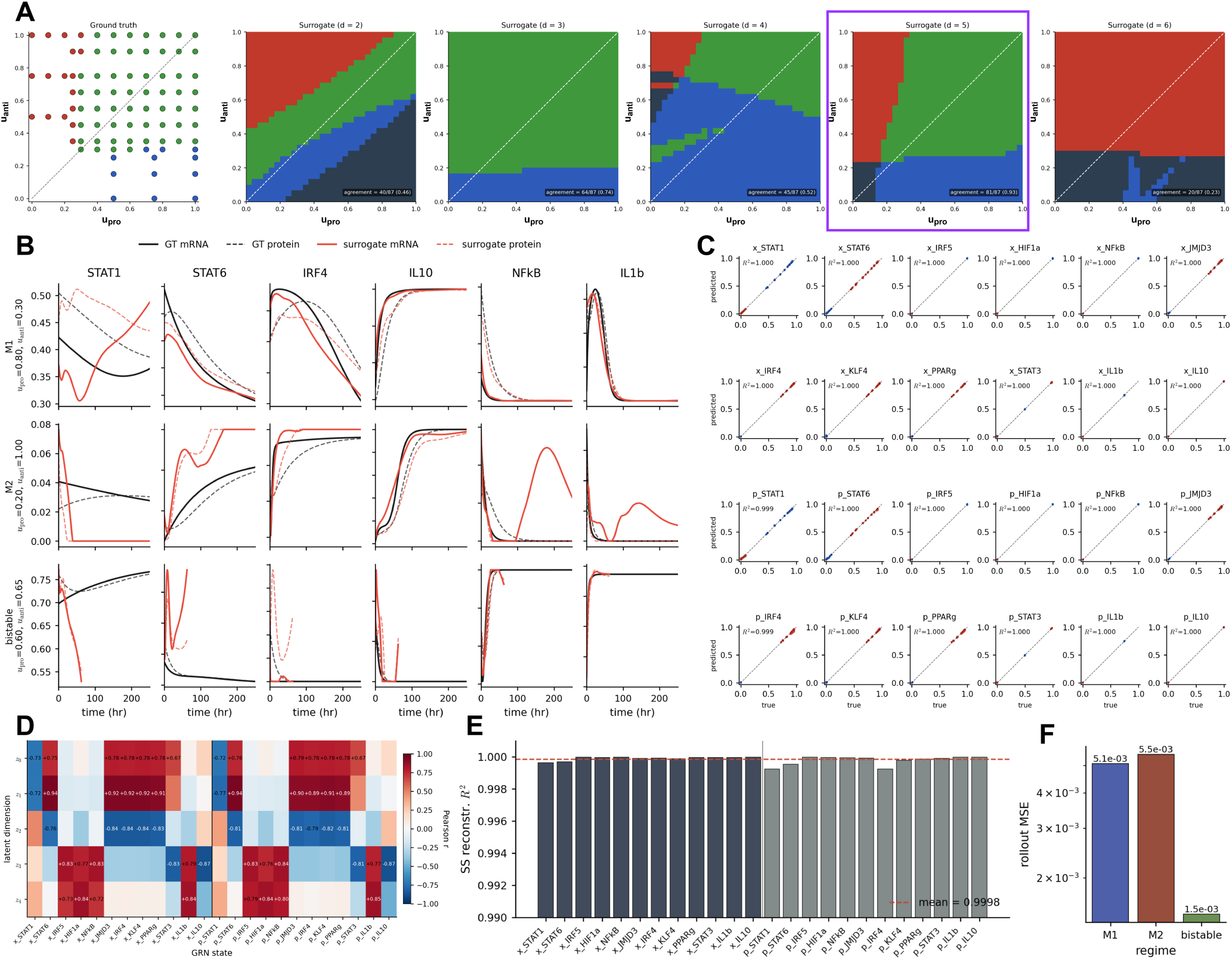
Autoencoder reconstruction fidelity and bistability preservation across latent dimensionalities. **(A)** Bifurcation regime classification on a 30×30 grid over (*u*_pro_, *u*_anti_) ∈ [0, 1]^2^ for the ground-truth GRN and surrogate models at *d* = 2-6. Colors denote monostable M1 (red), monostable M2 (blue), and bistable (green) regimes. The *d* = 5 surrogate (purple border) achieves 93.1% region agreement (81/87 classified input conditions), recovering all three dynamical regimes with boundary errors confined to the M1/M2 transition zone. **(B)** Trajectory rollout comparison between ground-truth GRN (black) and *d* = 5 surrogate (red) for six representative GRN states across M1 (*u*_pro_=0.80, *u*_anti_=0.30), M2 (*u*_pro_=0.20, *u*_anti_=1.00), and bistable (*u*_pro_=0.60, *u*_anti_=0.65) input regimes. Solid and dashed lines denote mRNA and protein respectively. Pearson correlations against ground-truth trajectories are 0.985, 0.991, and 0.996 in M1, M2, and bistable regimes, with MSEs of 5.1×10^−3^, 5.5×10^−3^, and 1.5×10^−3^ in normalized [0, 1]^24^ coordinates. **(C)** Steady-state reconstruction scatter plots for all 24 GRN states (12 mRNA, 12 protein) over 1344 stable steady states. All *R*^2^ values exceed 0.999, with the lowest observed for *p*_STAT1_ (*R*^2^ = 0.9993), *p*_STAT6_ (*R*^2^ = 0.9995), and *p*_IRF4_ (*R*^2^ = 0.9992). **(D)** Pearson correlation heatmap between the five latent dimensions *z*_0_-*z*_4_ and all 24 GRN states at steady state. *z*_1_ loads on STAT6 and the M2 cascade (JMJD3, IRF4, KLF4, PPAR*γ*; |*r*|>0.89); *z*_3_ and *z*_4_ co-represent the M1 inflammatory program; *z*_3_ captures the IL-10/STAT3 feedback axis (|*r*| >0.87) and *z*_4_ the HIF1*α*/IL-1*β*/IRF5 axis (|*r*| >0.84). **(E)** Per-state steady-state reconstruction *R*^2^ across all 24 GRN variables. The dashed red line marks the mean *R*^2^ = 0.9998. **(F)** Rollout MSE in normalized coordinates for M1, M2, and bistable input regimes. The lower bistable-regime error (1.5×10^−3^) reflects rapid attractor convergence suppressing long-time drift.

The ablation analysis further shows that the fixed-point pinning and phenotype separation losses are essential for preserving the multistable topology of the original GRN. Without *L*_fp_, the SINDy model is trained primarily to match latent velocities along trajectories and is not explicitly constrained to vanish at encoded steady states. As a result, the learned latent dynamics may reproduce transient motion while failing to place stable fixed points at the encoded attractor locations, degrading fixed-point continuation and bifurcation classification. Without *L*_sep_, the encoder can map M1-like and M2-like steady states into nearby or overlapping latent neighborhoods, especially near bistable boundaries where the two phenotypes may share similar aggregate expression levels. This collapse reduces the ability of the decoder and latent dynamics model to distinguish coexisting attractors, causing bistable regimes to be misclassified as monostable. Thus, *L*_fp_ and *L*_sep_ play complementary roles: *L*_fp_ anchors the learned dynamics at mechanistically known attractors, whereas *L*_sep_ enforces geometric separation between phenotypically distinct steady states.

### 3.2. Reconstruction and Bistability Preservation

Our trained *d* = 5 autoencoder achieves near-perfect steady-state reconstruction across all 24 GRN states, with a mean *R*^2^ = 0.9998 over 1344 stable steady states spanning the full (*u*_pro_, *u*_anti_) input space, as seen in Fig. 2C. All individual state *R*^2^ values exceed 0.999, with the lowest values observed for *p*_STAT1_ (*R*^2^ = 0.9993), *p*_STAT6_ (*R*^2^ = 0.9995), and *p*_IRF4_ (*R*^2^ = 0.9992), the three proteins most directly involved in the M1/M2 toggle, where the decoder must resolve sharp attractor separation.

Trajectory rollout fidelity is assessed by integrating the learned latent dynamics via RK4 from a single initial condition and decoding at each step. The surrogate achieves Pearson correlations of 0.985, 0.991, and 0.996 against ground-truth GRN trajectories in the M1, M2, and bistable input regimes, respectively, with corresponding mean-squared errors of 5.1 × 10^−3^, 5.5 × 10^−3^, and 1.5 × 10^−3^ (in normalized [0, 1]^24^ coordinates). The lower error in the bistable regime reflects trajectories that converge rapidly to a stable attractor, reducing long-time drift.

The bifurcation structure of the original 24-dimensional GRN is reproduced with 93.1% region agreement (81*/*87 input conditions) on a 30 × 30 grid over the (*u*_pro_, *u*_anti_) ∈ [0, 1]^2^ input space, illustrated in Fig. 2A. The surrogate correctly classifies monostable M1, monostable M2, and bistable input regimes by running latent fixed-point continuation from 200 initial conditions per grid point and decoding the converged attractors. The six misclassified points lie near the M1/M2 boundaries, where the latent fixed-point search is sensitive to the convergence tolerance.

Pearson correlation between the five latent coordinates and the 24 GRN states reveals a biologically interpretable decomposition, as shown in Fig. 2D. *z*_1_ is dominated by STAT6 and the downstream M2 cascade (JMJD3, IRF4, KLF4, PPARg; |*r*| > 0.89) . *z*_2_ captures the same M2 program with opposite sign (|*r*| > 0 83), encoding the nonlinear curvature of the M2 manifold not captured by *z*_1_ alone. *z*_3_ and *z*_4_ co-represent the M1-associated inflammatory program, with *z*_3_ loading on the IL-10/STAT3 feedback axis (|*r*| > 0.87) and *z*_4_ on the HIF1*α*/IL-1β/IRF5 axis (|*r*| > 0.84). *z*_0_ captures shared M2 effector activity (JMJD3, KLF4; |*r*| *≈* 0.79). The STAT1/STAT6 toggle is distributed across *z*_0_-*z*_2_, consistent with the mutual repression topology precluding a single linear axis from separating the two attractors.

Finally, we examine the evolution of an initially unimodal macrophage population under a bifurcating perturbation, with (*u*_pro_ = *u*_anti_ = 0.9). Fig. 3M shows the resulting particle cloud in latent space from multiple two-dimensional projections, such as (*z*_1_) − (*z*_2_), (*z*_2_) − (*z*_3_), and other coordinate pairs, over the interval *t* = 0 to *t* = 1000 h. This relatively long simulation window was chosen to allow the system to approach equilibrium, particularly because slow components of the GRN, such as STAT1 mRNA with a decay rate of 0.0067 h^−1^, introduce long relaxation timescales. The latent coordinates (*z*_1_, *z*_2_, *z*_3_, …) should not be interpreted as individual biological variables. For example, a change from *z*_1_ = 0.5 to *z*_1_ = 1 does not correspond directly to a specific molecular or phenotypic change in the cell. Rather, the latent space provides a reduced representation in which the organization and evolution of the macrophage population can be visualized.

**Figure 3.**
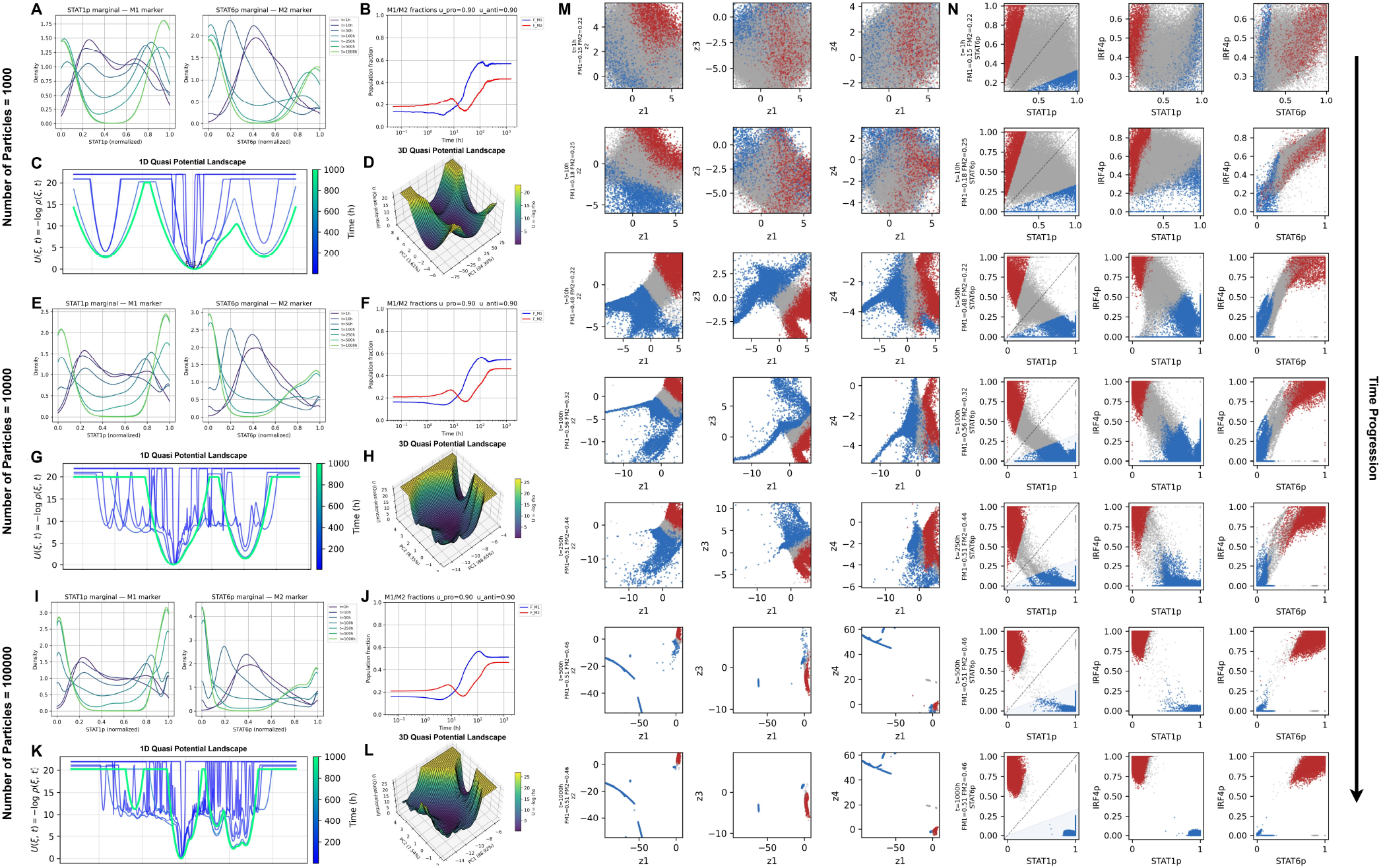
Population-level bistability recovered by the latent particle solver. **(A-D)** Results for *N*_*p*_ = 10^3^ particles. Kernel density estimates of reconstructed *p*_STAT1_ and *p*_STAT6_ show that an initially unimodal population evolves into a bimodal distribution over *t* = 0-1000 h, consistent with separation into M1-like and M2-like phenotypic subpopulations. The corresponding phenotype fractions show the emergence of both classified populations from an initially unresolved ensemble: blue denotes the M1-like fraction *F*_*M*1_, red denotes the M2-like fraction *F*_*M*2_, and gray denotes mixed or unclassified particles. Under equal pro- and anti-inflammatory cytokine forcing, the M1-like and M2-like fractions approach comparable values at long times. The associated one-dimensional and three-dimensional quasi-potential landscapes visualize the formation of two phenotypic basins separated by a barrier, following the landscape construction for multistable systems (Zhou et al., 2012). **(E-H)** and **(I-L)** The same diagnostics for *N*_*p*_ = 10^4^ and *N*_*p*_ = 10^5^ particles, respectively. Increasing *N*_*p*_ preserves the bimodal marker distributions, the two-basin quasipotential landscape, and the long-time M1/M2 population fractions, indicating convergence of the particle approximation. **(M)** Time evolution of the particle ensemble in the learned latent space, shown through three two-dimensional coordinate projections. Each row corresponds to a later time point, and each column gives a different projection of the same latent particle cloud. Particles are colored by their decoded phenotype classification: M1-like particles are blue, M2-like particles are red, and mixed or unresolved particles are gray. The initially unresolved cloud progressively separates into two phenotypically distinct clusters, demonstrating that the latent dynamics organize the population into separate attractor basins. **(N)** The same particle ensemble decoded back into the original 24-dimensional GRN state space and projected onto representative physical coordinates. The decoded population separates into distinct blue M1-like and red M2-like subclusters, confirming that the PBE solved in latent space, when propagated back through the decoder, preserves the bistable phenotypic structure of the full GRN.

At *t* = 0, most particles appear as a large gray cloud of mixed or unresolved macrophage states. This is expected because the external stimulus has just been applied, and the downstream mRNA and protein levels have not yet accumulated sufficiently for the phenotype classifier to assign a clear M1-like or M2-like identity. As the system evolves, particles are classified according to the relative activation of M1- and M2-associated GRN modules. Specifically, the M1-like module score is computed from the protein levels of STAT1, IRF5, HIF1*α*, NF*κ*B, and IL1*β*, while the M2-like module score is computed from STAT6, JMJD3, IRF4, KLF4, PPAR*γ*, STAT3, and IL10. Geometric-mean scores, denoted by *S*_*M*1_ and *S*_*M*2_, are used so that classification reflects coordinated activation of the broader regulatory module rather than the dominance of a single marker (Haynes, Vallania, Liu, Bongen, Tomczak, Andres-Terrè, Lofgren, Tam, Deisseroth, Li et al., 2017; Chawla, Cappuccio, Tamminga, Sealfon, Zaslavsky and Kleinstein, 2022). A signed polarization index is then defined as

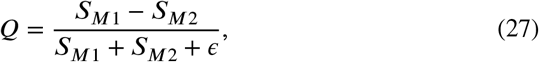

where (*Q* > 0) indicates an M1-like bias and (*Q* < 0) indicates an M2-like bias. Particles are classified as M1-like or M2-like only when (|Q|) exceeds a prescribed polarization threshold and the corresponding module score exceeds a minimum activation threshold. Otherwise, they are labeled as mixed or unresolved. Notice how by *t* = 1000 h, the initially unresolved gray cloud has separated into distinct blue and red subclusters, indicating that the macrophage population has split into divergent regions of the latent space corresponding to M1-like and M2-like phenotypic programs. This split is also confirmed by the population fractions and the number-density plots for STAT1p and STAT6p that morph from unimodal to bimodal over time.

In vivo, individual cells are continuously exposed to a complex cytokine environment, with IFN-*γ* and IL-4 (primarily secreted by T-cells) serving as two of the dominant opposing polarization cues. Together with other cytokines, these signals strongly influence macrophage polarization dynamics. As shown in Fig. 1A, the mechanistic GRN considered in this work will always exhibit bistable behavior, allowing the system to evolve toward either M1- or M2-like phenotypic states depending on the relative strength of the external stimuli. From a dynamical systems perspective, such bistability is particularly important in chronic inflammatory conditions, where macrophages may become trapped within persistent pathological attractors due to dysregulated signaling or aberrant gene expression (Geiß, Salas, Guevara-Coto, Régnier-Vigouroux and Mora-Rodríguez, 2022). Consequently, understanding how specific parameters alter the attractor structure and basin geometry may provide mechanistic insight into how macrophage populations can be redirected away from chronic inflammatory states.

### 3.3. Hysteresis under Time-varying Inputs

In the original GRN state space, the cytokine-dependent dynamics is written abstractly as 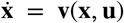, where x ∈ ℝ^24^ denotes the full GRN state and **u** denotes the external cytokine inputs. Sweeping *u*_pro_ at fixed *u*_anti_ reveals a bistable region in which two stable equilibria coexist, corresponding broadly to M1-like and M2-like phenotypic states. The forward and backward sweeps follow different equilibrium branches, producing a hysteresis loop (see Fig. 1B). The measured loop width 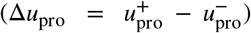, therefore quantifies the range of *u*_pro_ values over which bistability persists. Here, 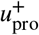 and 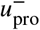 denote the upper and lower transition thresholds respectively. The bistable region consists of two discrete stable equilibria separated by an unstable saddle. As *u*_pro_ increases, the system remains on the branch associated with its current basin of attraction until the corresponding stable equilibrium is lost, typically through a saddle-node bifurcation. At that point, the trajectory jumps to the remaining stable branch. When *u*_pro_ is subsequently decreased, the system does not retrace the same path because it is now initialized on the opposite branch. It remains there until the second saddle-node threshold is crossed. Thus, the closed hysteresis loop arises from the coexistence of two stable phenotypic attractors and the separation of their transition thresholds (Chow, Morris and Rabbah, 2023).

This interpretation extends naturally to the particle formulation (Fig. 4). Rather than tracking a single GRN trajectory, the latent-space PBE describes the evolution of a population density, ñ(**z**, *t*), over the learned latent coordinates. Therefore, the observed hysteresis is not merely a property of one initialized trajectory, but of the evolving particle ensemble. In Figure 4, the particle cloud does not retrace the same path when the cytokine schedule is reversed. On the forward path, particles initially associated with the M2-like branch remain there until *u*_pro_ crosses the upper threshold 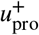, at which point the M2-like attractor is lost and the ensemble transitions toward the M1-like branch. On the return path, the ensemble is already M1-committed and remains near that branch until *u*_pro_ crosses the lower threshold 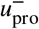. Since 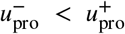, the forward and return paths occupy different regions of latent space, producing the observed hysteretic behavior.

**Figure 4.**
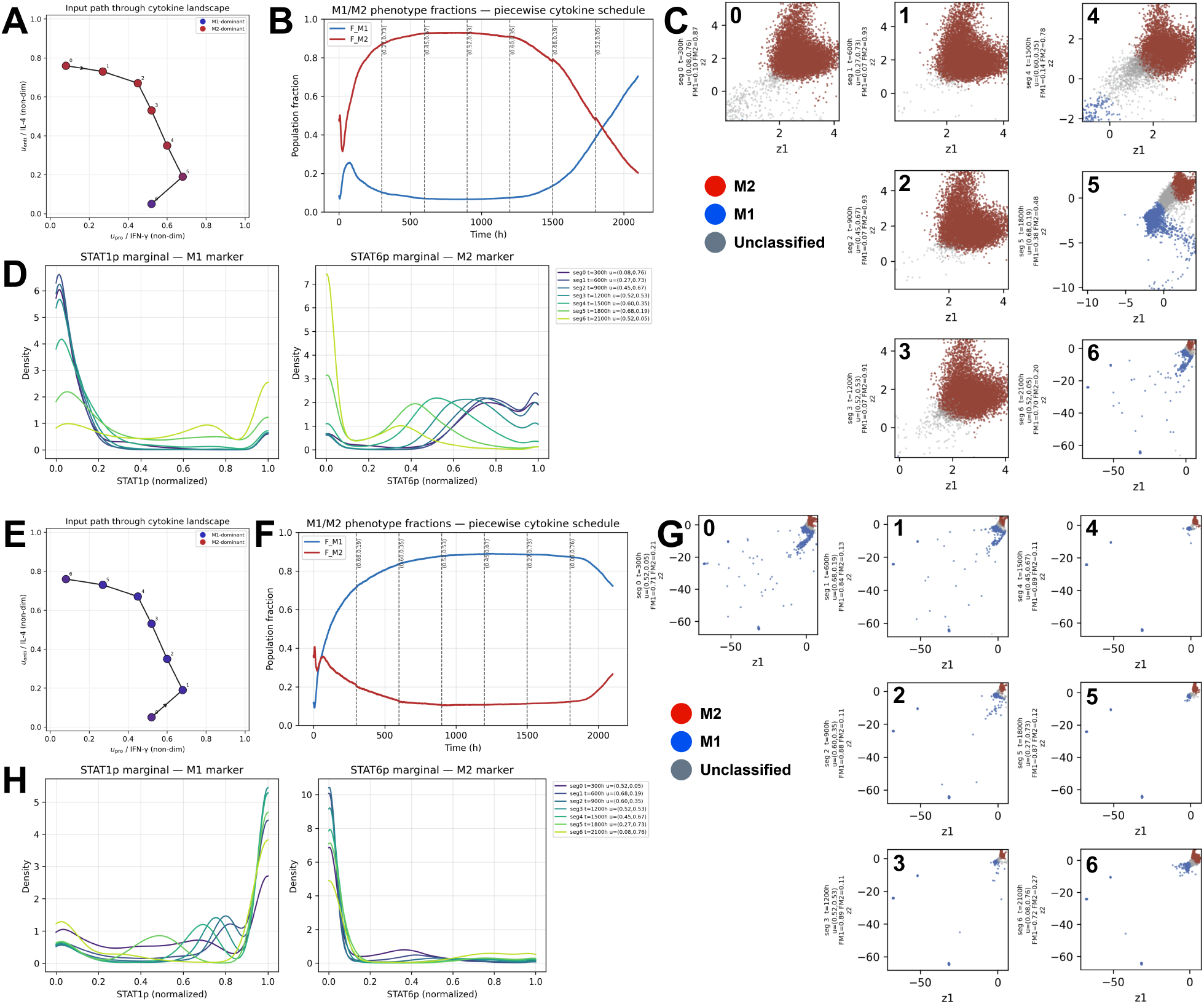
Population-level hysteresis under piecewise-constant cytokine schedules. **(A-D)** Forward M2-to-M1 schedule. **(A)** Input path in the cytokine landscape, with *u*_pro_ increased relative to *u*_anti_. **(B)** Time evolution of the decoded phenotype fractions. The population initially remains predominantly M2-like, then transitions toward an M1-like state as the forward switching threshold is crossed. Blue denotes the M1-like fraction and red denotes the M2-like fraction. **(C)** Latent-space particle snapshots along the forward schedule, projected onto representative latent coordinates and colored by decoded phenotype: blue, M1-like; red, M2-like; gray, mixed or unresolved. The particle cloud remains organized near the M2-like region before shifting toward the M1-like branch. **(D)** Reconstructed *p*_STAT1_ and *p*_STAT6_ marginal distributions along the same schedule, showing the corresponding marker-level transition from M2-like to M1-like polarization. **(E-H)** Reverse M1-to-M2 schedule. **(E)** Reversed cytokine path through the same input landscape. **(F)** Phenotype fractions during the return sweep, showing that the population remains M1-like over a different range of cytokine values before switching back toward the M2-like state. **(G)** Latent-space particle snapshots along the reverse schedule. At comparable cytokine values, the cloud occupies a different region of latent space than in the forward sweep, reflecting memory of the previously occupied attractor basin. **(H)** Reconstructed *p*_STAT1_ and *p*_STAT6_ marginals during the reverse schedule, confirming the delayed return from M1-like to M2-like polarization. Together, the forward and reverse schedules demonstrate population-level hysteresis: the decoded phenotype distribution and latent particle cloud depend not only on the instantaneous cytokine input, but also on the direction of the sweep and the attractor branch occupied beforehand.

This explains why the latent particle clouds differ between the M2-to-M1 and M1-to-M2 cytokine schedules. Although the two schedules pass through comparable regions of cytokine-input space, the particle cloud does not depend only on the instantaneous value of **u**(*t*). It also depends on the branch occupied immediately before that input value was reached. During the M2-to-M1 schedule, particles remain organized around the M2-like attractor until the upper transition threshold is crossed. During the reverse M1-to-M2 schedule, the particles begin from the M1-like branch and remain there until the lower transition threshold is crossed. Thus, at intermediate cytokine values, the latent-space cloud can occupy different regions depending on the direction of the sweep. This path dependence is the population-level signature of hysteresis.

This behavior is visible in Fig 4C and 4G: the particle clouds generated during the M2-to-M1 schedule and the M1-to-M2 schedule do not retrace one another in latent space. Instead, the ensemble retains memory of its previous phenotypic branch, producing distinct cloud geometries and different M1/M2 phenotype fractions along the forward and reverse paths. The path-dependent separation is strongest when the cytokine path crosses the bistable region of the GRN. Outside this region, where the system is effectively monostable, the forward and reverse trajectories are expected to collapse toward the same attracting branch.

An important feature of the present implementation is that cytokine inputs are applied as piecewise-constant step functions. This choice preserves a fixed encoder geometry within each time interval. For a scalar function *f* (x_*t*_, *t*), Itô’s lemma (Chiarella, He and Nikitopoulos, 2015) gives

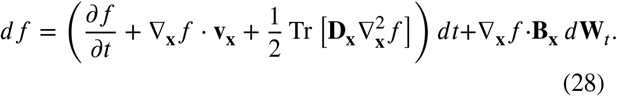

In our model, the encoder is trained as a map **z** = *ε*(x), and does not take **u** as an explicit input. If the input were absorbed directly into the encoder as *ε* (x, **u**), then a continuously varying input would introduce the additional contribution 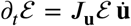 which would add a term involving 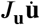 to the latent drift. To avoid this issue, the cytokine schedule is imposed as

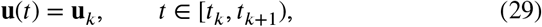

so that *d***u***/dt* = 0 almost everywhere within each segment. Consequently, within each interval, the encoder has no explicit time dependence. At each jump time *t*_*k*_, the drift field changes instantaneously from **v**(⋅, **u**_*k*−1_) to **v**(⋅, **u**_*k*_), but the particle positions remain continuous. When the input changes from **u**_*k*_ to **u**_*k*+1_, the attractor locations and basin boundaries shift. Hysteresis emerges because the particle ensemble carries memory of its previous branch: on the forward sweep, particles transition only after the M2-like basin is lost, whereas on the reverse sweep, particles remain in the M1-like basin until the lower transition threshold is reached. Therefore, the latent-space hysteresis loop reflects the same bistable basin structure observed in the original GRN state space.

### 3.4. Simulating Gene Knockout Perturbations

Gene knockout perturbations provide a useful way to test whether the reduced latent-space framework preserves the expected functional consequences of specific regulatory disruptions. In the present study, we simulate three knockout conditions: IRF4 knockout (*Irf* 4^−*/*−^), IRF5 knockout (*Irf* 5^−*/*−^), and a double IRF4 + IRF5 knockout (*Irf* 4^−*/*−^+ *Irf* 5^−*/*−^). These perturbations are particularly informative because IRF4 and IRF5 occupy opposing positions in the macrophage polarization network. IRF4 is associated with anti-inflammatory and tissue-repairing M2-like polarization, whereas IRF5 is a key regulator of pro-inflammatory M1-like activation. Therefore, selectively removing either transcription factor provides a direct test of whether the coupled autoencoder-particle solver can reproduce the expected shift in phenotypic balance.

In the GRN considered here, IRF4 forms part of the STAT6-driven regulatory cascade that promotes the M2-like program. Knocking out IRF4 therefore weakens this anti-inflammatory branch (Eguchi, Kong, Tenta, Wang, Kang and Rosen, 2013). However, the M2 program is not expected to vanish completely, since STAT6 can still directly regulate some downstream M2-associated targets. At the same time, IRF4 inhibits components of the M1-associated branch, including IRF5 and NF-(*κ*)B. Removing IRF4, therefore, releases this inhibition and allows the pro-inflammatory module to become more strongly activated. As a result, the IRF4 knockout is expected to shift the population toward an M1-like phenotype. Conversely, IRF5 promotes the M1-like program and supports inflammatory cytokine production. Knocking out IRF5 suppresses this pro-inflammatory branch and is therefore expected to bias the population away from M1-like polarization and toward an M2-like phenotype (Petrova, Sherstyukova, Kandrashina, Inozemtsev, Tsitrina, Fedorova, Shvedov, Kuzovlev, Dokukin, Kotelevtsev et al., 2026). The double IRF4 + IRF5 knockout removes regulatory support for both opposing branches, making it a useful perturbation for examining how the network behaves when both major polarization-associated transcriptional regulators are disrupted.

Figure 5 summarizes the simulated knockout responses under the bifurcating stimulation condition (*u*_pro_ = *u*_anti_ = 0.9). For each knockout, we show the marginal distributions of STAT1p and STAT6p as representative M1- and M2-associated markers, the time evolution of the GRN-based phenotype fractions, the signed polarization index (Q), and the corresponding latent-space particle clouds. In the IRF4 knockout case, the population rapidly loses M2-like support and evolves toward a predominantly M1-like state. This is reflected by the increasing M1-like phenotype fraction, the positive shift in the polarization index (Q), and the organization of the latent particle cloud into a region dominated by M1-like particles. This behavior is consistent with the GRN topology: removal of IRF4 both weakens the STAT6-IRF4-STAT3 axis and relieves inhibition of the M1-associated regulatory module (Fig. 5A-5E).

**Figure 5.**
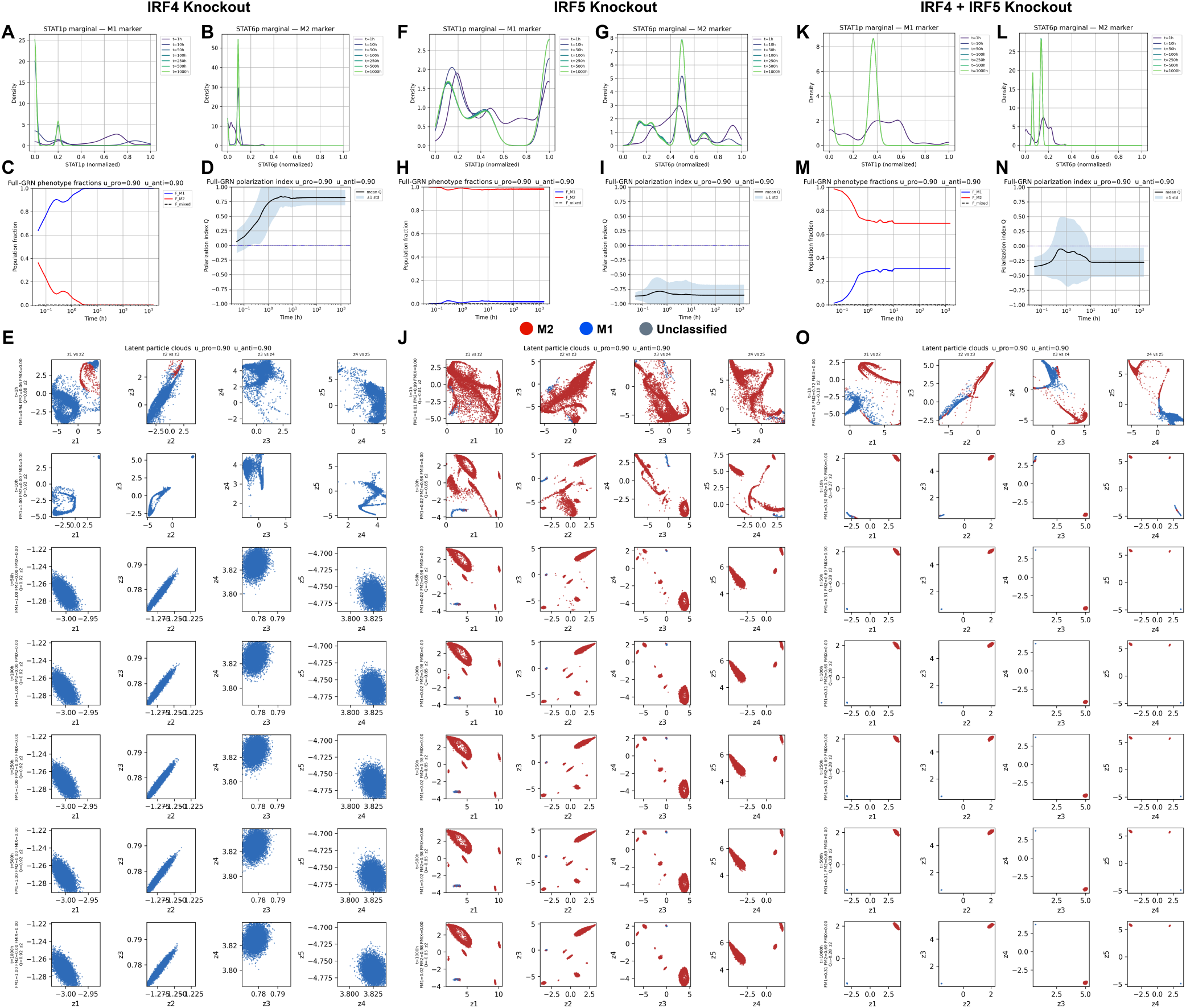
Population-level response to transcription-factor knockout perturbations. **(A-E)** IRF4 knockout. Because IRF4 participates in the STAT6-driven M2-like regulatory cascade and inhibits components of the M1-associated branch, its removal weakens anti-inflammatory support and releases pro-inflammatory activation. Across the reconstructed *p*_STAT1_ and *p*_STAT6_ marginal distributions, phenotype fractions, polarization index *Q*, and latent particle projections, the population shifts toward an M1-like state. Blue denotes M1-like particles, red denotes M2-like particles, and gray denotes mixed or unresolved particles. **(F-J)** IRF5 knockout. Because IRF5 promotes the M1-like inflammatory program, its removal suppresses pro-inflammatory polarization and biases the population toward an M2-like response. This is reflected by weakened *p*_STAT1_ activation, increased M2-like phenotype fraction, a negative shift in the polarization index *Q*, and latent particle clouds dominated by red M2-like particles. **(K-O)** Double IRF4 + IRF5 knockout. Simultaneous removal of both opposing transcriptional regulators produces a more restricted and network-dependent polarization response. Rather than reproducing either single-knockout phenotype, the particle ensemble exhibits reduced access to the full M1-M2 polarization structure, with the final phenotype distribution determined by the remaining GRN topology and the external cytokine inputs. Across all perturbations, the decoded marker distributions, phenotype fractions, polarization indices, and latent particle organization recover the expected qualitative shifts in macrophage polarization, indicating that the reduced latent-space PBE framework preserves biologically relevant responses to targeted gene knockouts.

In contrast, the IRF5 knockout produces the opposite qualitative response. Since IRF5 is a major driver of the M1-like program, its removal suppresses pro-inflammatory polarization and shifts the population toward an M2-like state. This is seen in the negative polarization index and the predominance of M2-like particles in the latent projections. The STAT1p marginal distribution is also weakened relative to the IRF4 knockout case, consistent with reduced activation of the M1-associated branch. Thus, the latent-space particle solver correctly captures the expected phenotypic consequence of removing a pro-inflammatory transcriptional regulator (Fig. 5F-5J).

The double IRF4 + IRF5 knockout produces a more constrained response because both major opposing regulators are removed simultaneously. Rather than producing the same strong polarization observed in the single-knockout cases, the population exhibits a more limited separation in latent space, with reduced access to the full M1-M2 polarization structure. The resulting phenotype distribution reflects the residual asymmetry of the remaining GRN: although both branches are disrupted, downstream and parallel regulatory interactions can still support partial activation of one program over the other. This case is important because it shows that the framework does not simply assign phenotypes based on the deleted genes alone; instead, the final population-level behavior emerges from the remaining network topology, the external cytokine inputs, and the dynamical evolution of the particle ensemble (Fig. 5K-5O).

Overall, these knockout simulations provide a mechanistic consistency check for the coupled autoencoder-particle solver. The IRF4 knockout shifts the system toward an M1-like response, the IRF5 knockout shifts it toward an M2-like response, and the double knockout produces a more restricted and network-dependent polarization outcome. Importantly, these qualitative behaviors are recovered not only in the original GRN variables, but also in the latent-space particle representation, indicating that the reduced framework preserves biologically relevant responses to targeted genetic perturbations.

## 4. Discussion

In this work, we constructed a dynamics-preserving autoencoder to enable the solution of a high-dimensional population balance equation (PBE) in a reduced latent space. Although the present study focuses on PBEs arising from cell-state dynamics, the underlying idea is not restricted to this specific class of equations. In principle, similar reductions can be developed for other high-dimensional linear PDEs, provided that the transformation to latent space preserves the relevant dynamical structure of the original system. Extensions to quasilinear or nonlinear PDEs are also possible, but such applications would require additional mathematical structure to ensure that the reduced representation remains faithful to the original dynamics. The motivation for this approach comes from the inherently high-dimensional nature of cellular systems. A realistic cell-state model may involve many genes, proteins, metabolites, and regulatory interactions, making the governing dynamics difficult to solve directly. A reduced representation is therefore necessary, but the reduction must simplify the computation without destroying the essential physics and regulatory structure of the original network. This is especially important in the context of single-cell data, where modern measurements resolve cell populations at the distributional level. Such data motivates models that can describe not only an average cellular state, but the evolution of an entire distribution of heterogeneous cell states.

Macrophage polarization was chosen as the biological system of interest for demonstrating this framework because it is a crucial process across many inflammatory and disease contexts, including wound healing, chronic inflammation, and tumor-associated macrophage behavior in cancer (Chen, Saeed, Liu, Jiang, Xu, Xiao, Rao and Duo, 2023c). In these settings, macrophage phenotype is not fixed. Rather, cells must transition between inflammatory, resolving, and tissue-repairing programs in response to changing environmental signals. When this transition occurs at the wrong time, with the wrong magnitude, or becomes trapped in a persistent state, the resulting imbalance can disrupt tissue repair and contribute to pathological inflammation (Pomeyie, Abrokwah, Boison, Amoani, Kyei, Adinortey and Barnie, 2025).

At the molecular level, such failures are often associated with dysregulated gene expression, altered signaling activity, or misregulation of key transcription factors and cytokine-responsive pathways. Experimentally, these mechanisms are commonly investigated using perturbations such as gene knockouts, knockdowns, or pathway inhibition. Here, we show that the population-level consequences of such regulatory perturbations can also be studied in silico using the coupled autoencoder-particle solver framework. Although exact quantitative prediction requires experimental calibration and validation, the model can still provide useful qualitative insight into the direction and structure of the population response. For example, it can predict whether a perturbation shifts the macrophage population toward an M1-like or M2-like region, whether it broadens or collapses the phenotypic distribution, and whether it alters the accessibility of different attractor states. Such predictions can help guide experimental design by identifying which perturbations are most likely to produce informative or biologically meaningful outcomes.

More broadly, this work points toward a future in which increasingly large regulatory networks can be embedded into reduced, dynamics-preserving spaces and used to simulate distribution-level cellular responses. While the present study focuses on a simplified macrophage GRN, the long-term goal is to move toward richer, data-informed regulatory representations that can connect single-cell measurements, mechanistic network models, and predictive perturbation analysis.

## CRediT authorship contribution statement

**Prateek Gupta:** Conceptualization, Data curation, Formal analysis, Investigation, Methodology, Software, Validation, Visualization, Writing - original draft, Writing - review & editing; **Shourya Verma**: Conceptualization, Data curation, Formal analysis, Investigation, Methodology, Software, Validation, Visualization, Writing - original draft, Writing - review & editing; **Ananth Grama**: Project administration, Methodology, Resources, Supervision, Writing - review & editing; **Doraiswami Ramkrishna**: Conceptualization, Investigation, Methodology, Project administration, Resources, Supervision, Writing - review & editing.

## Data Availability Statement

All model-generated data used in this study were produced using Python scripts developed by the authors. The codes used to generate, process, and analyze these data are curated on the following GitHub repository: https://github.com/shouryaverma/HighDimPBE. Parameter values and experimental quantities used for model construction were obtained from publicly available sources, including TTDB and CHX-chase measurements, and are reported in the main text and Supplementary Information.

## Acknowledgments

The authors would like to acknowledge helpful discussions with Mengbo Wang (Purdue Computer Science). Generative AI tools (OpenAI’s ChatGPT and Anthropic’s Claude) were used to assist with grammar correction, language refinement, and manuscript editing. The authors reviewed and edited all AI-assisted text. No scientific conclusions, analyses, results, or interpretations were generated by these tools. The authors take full responsibility for the accuracy and integrity of the manuscript.

## Footnotes

1 ñ ≠ n/|J_ε_ |; J_ε_ is not invertible.

