## Supplemental File S1 for "Solving High-Dimensional Population Balance Equations via Dynamics-Preserving Autoencoders"

### Supplementary File S1

Prateek Gupta, Shourya Verma, Ananth Grama, Doraiswami Ramkrishna

June 2026

### Contents

**1 Full GRN Model Specification** **1**

**2 Diffusion Tensor** **2**

**3 Full SINDy library construction and discovered equations** **4**

**4 PBE transformation under autoencoder** **6**

**5 Latent-space stochastic particle update** **7**

**6 Training losses** **7**

### 1 Full GRN Model Specification

The network models 12 genes, yielding 24 state variables for mRNA and protein. The indices correspond to the following genes: 1: STAT1, 2: STAT6, 3: IRF5, 4: HIF1$\alpha$, 5: NF$\kappa$B, 6: JMJD3, 7: IRF4, 8: KLF4, 9: PPAR$\gamma$, 10: STAT3, 11: IL-1$\beta$, 12: IL-10. To simplify the notation, we define the Hill activation $A_i(p_i)$, Hill repression $R_i(p_i)$, and external input activation $A_{in}(u)$ functions as:

$$A_i(p_i) = \frac{p_i^{n_H}}{K_i^{n_H} + p_i^{n_H}}, \quad R_i(p_i) = \frac{K_i^{n_H}}{K_i^{n_H} + p_i^{n_H}}, \quad A_{in}(u) = \frac{u^{n_H}}{0.5^{n_H} + u^{n_H}} \tag{1}$$

where $n_H = 3$ is the Hill coefficient and $K_i$ is the Hill threshold for gene $i$. The dynamics for the 12 mRNA species are given by:

$$\frac{dm_1}{dt} = \alpha_1 + \gamma_1\big(w_u A_{in}(u_{pro}) + w_s A_1(p_1)\big)R_2(p_2) - \delta_1 m_1 \tag{2a}$$

$$\frac{dm_2}{dt} = \alpha_2 + \gamma_2\big(w_u A_{in}(u_{anti}) + w_s A_2(p_2)\big)R_1(p_1) - \delta_2 m_2 \tag{2b}$$

$$\frac{dm_3}{dt} = \alpha_3 + \gamma_3 A_5(p_5)R_7(p_7)R_{10}(p_{10}) - \delta_3 m_3 \tag{2c}$$

$$\frac{dm_4}{dt} = \alpha_4 + \gamma_4 A_5(p_5) - \delta_4 m_4 \tag{2d}$$

$$\frac{dm_5}{dt} = \alpha_5 + \gamma_5 \Phi_{act}\Phi_{rep} - \delta_5 m_5 \tag{2e}$$

$$\frac{dm_6}{dt} = \alpha_6 + \gamma_6 A_2(p_2) - \delta_6 m_6 \tag{2f}$$

$$\frac{dm_7}{dt} = \alpha_7 + \gamma_7 A_2(p_2)\big(0.05 + 0.95A_6(p_6)\big) - \delta_7 m_7 \tag{2g}$$

$$\frac{dm_8}{dt} = \alpha_8 + \gamma_8 A_2(p_2)\big(0.5 + 0.5A_7(p_7)\big) - \delta_8 m_8 \tag{2h}$$

$$\frac{dm_9}{dt} = \alpha_9 + \gamma_9 A_2(p_2)\big(0.5 + 0.5A_8(p_8)\big) - \delta_9 m_9 \tag{2i}$$

$$\frac{dm_{10}}{dt} = \alpha_{10} + \gamma_{10} A_{12}(p_{12})\big(0.5 + 0.5A_9(p_9)\big) R_3(p_3) - \delta_{10} m_{10} \tag{2j}$$

$$\frac{dm_{11}}{dt} = \alpha_{11} + \gamma_{11} A_5(p_5)\big(0.5 + 0.5A_4(p_4)\big) - \delta_{11} m_{11} \tag{2k}$$

$$\frac{dm_{12}}{dt} = \alpha_{12} + \gamma_{12} A_{10}(p_{10}) - \delta_{12} m_{12} \tag{2l}$$

where the NF$\kappa$B ($i = 5$) activation logic acts as an OR-gate among STAT1, IL-1$\beta$, and IRF5, while its repression acts as an AND-gate among the M2 cascade members:

$$\Phi_{act} = 1 - \big(1 - A_1(p_1)\big)\big(1 - A_{11}(p_{11})\big)\big(1 - A_3(p_3)\big)$$
$$\Phi_{rep} = R_2(p_2) R_7(p_7) R_8(p_8) R_9(p_9) R_{10}(p_{10})$$

Because the system is solved without assuming quasi-steady state (QSS) for translation, the remaining 12 ordinary differential equations govern the exact protein dynamics. For each gene $i \in \{1, \ldots, 12\}$, the protein evolution is:

$$\frac{dp_i}{dt} = \beta_i x_i - \mu_i p_i \tag{3}$$

where $\mu_i$ is the protein decay rate. To ensure the biological fidelity of the model, kinetic parameters were anchored to experimentally determined ranges where possible. The mRNA decay rates ($\delta_i$) for all 12 network genes were sourced directly from the Transcriptome-wide Turnover Database (TTDB), while protein degradation rates ($\mu_i$) were derived from cycloheximide (CHX) chase assays [1 - 7]. The gene-specific parameters are detailed in Table 1, while the shared global network parameters are listed in Table 2.

For the transcriptional kinetics, we assume that the Hill exponent ($n_H = 3$) does not represent the physical cooperativity typically observed in DNA-transcription factor binding events. Instead, we utilize $n_H$ as an ultra-sensitivity coefficient that allows the regulatory genes to transition between 'ON' and 'OFF' states in a switch-like manner [8].

The basal transcription rates ($\alpha_i$) were estimated based on global transcriptomics. In murine macrophages, the total transcript production is on the order of $10^4 - 10^5$ transcripts per cell per hour, corresponding to a genome-wide average of $0.01 - 3$ transcripts per gene per hour. However, 40–50% of the cellular genes are housekeeping genes [9], which account for approximately 90% of the total transcript pool and are $100 - 1000$ times more abundant than transcription factor (TF) genes. The 12 genes comprising our modeled GRN (e.g., STAT1, NF$\kappa$B) are almost exclusively TFs and receptor proteins. With typical unstimulated expression levels of $1 - 50$ FPKM (compared to GAPDH at $\sim 1000 - 5000$ FPKM), these network genes sit in the bottom 20–30% of the expression distribution. Consequently, their basal transcription rates are roughly $10 - 100$ times below the genome-wide average. Therefore, we set the basal transcription rate uniformly for the GRN to $\alpha = 0.005$ transcripts/(gene-hour), well within the biologically justified range of $0.001 - 0.1$ for TF and receptor genes.

Table 1: Gene-specific dimensional kinetic parameters.

| **Gene** | **mRNA Decay, $\delta_i$ (hr$^{-1}$)** | **Protein Decay, $\mu_i$ (hr$^{-1}$)** | **Translation, $\beta_i$ (hr$^{-1}$)** |
|---|---|---|---|
| STAT1 | 0.0067 | 0.029 | 0.15 |
| STAT6 | 0.044 | 0.099 | 0.50 |
| IRF5 | 0.30 | 0.17 | 0.15 |
| HIF1$\alpha$ | 0.47 | 1.00 | 0.20 |
| NF$\kappa$B | 0.33 | 0.154 | 0.20 |
| JMJD3 | 1.08 | 0.17 | 0.30 |
| IRF4 | 2.96 | 0.077 | 0.40 |
| KLF4 | 0.74 | 0.347 | 0.30 |
| PPAR$\gamma$ | 0.25 | 0.365 | 0.25 |
| STAT3 | 0.26 | 0.063 | 0.15 |
| IL-1$\beta$ | 0.87 | 0.693 | 0.50 |
| IL-10 | 0.33 | 0.193 | 0.20 |

### 2 Diffusion Tensor

A mechanistic diffusion kernel $\mathbf{D} \in \mathbb{R}^{24 \times 24}$ can be specified using a coarse-grained approximation. In the context of a discrete Chemical Master Equation (CME) or a Chemical Langevin Equation (CLE), one can define a system of $R$ elementary

Table 2: Shared regulatory parameters governing the toggle core and downstream cascades.

| Parameter | Description | Value | Units |
|---|---|---|---|
| $K_p$ | Absolute Hill threshold for protein activation | 1200.0 | nM |
| $n$ | Hill coefficient (cooperativity) | 3 | – |
| $\alpha$ | Basal transcription rate (uniform for all genes) | 0.005 | $\mathrm{hr}^{-1}$ |
| $w_u$ | Additive toggle weight for external input | 0.5 | – |
| $w_s$ | Additive toggle weight for self-excitation | 0.5 | – |
| $\gamma_1$ | Max transcription rate for STAT1 | 5.0 | $\mathrm{hr}^{-1}$ |
| $\gamma_6$ | Max transcription rate for STAT6 ($\gamma_1 \cdot \delta_2/\delta_1$) | 32.836 | $\mathrm{hr}^{-1}$ |
| $\gamma_{downstream}$ | Downstream transcription multiplier ($5 \cdot \delta_i \cdot K_p$) | $6000 \cdot \delta_i$ | $\mathrm{hr}^{-1}$ |

reactions involving $N$ species such that [10]:

$$\sum_{i=1}^{N} s_{ir}\mathbf{x}_i \xrightarrow{k_r} \sum_{i=1}^{N} s'_{ir}\mathbf{x}_i \quad \forall\ r \in \{1, ..., R\} \tag{4}$$

and for which, a transition rate (or a propensity) $p_r$ can be defined such that, given the state of the system is $\mathbf{x}$ at time $t$, the probability of the $r^{th}$ transition occurring (or of the $r^{th}$ reaction firing) in the infinitesimal time interval $[t, t + dt)$ is:

$$P(x + \nu_r, t + dt|x, t) = p_r dt + o(dt) \tag{5}$$

where $\nu_r = \mathbf{s}'_r - \mathbf{s}_r$ is the stoichiometric vector and $o(dt)$ denotes terms that vanish faster than $dt$ as $dt \to 0$. Subsequent Kramer-Moyal (KM) expansion of the CME and second-order truncation of this expansion then gives a continuous Fokker-Planck Equation (FPE) which can be written as:

$$\frac{\partial \rho}{\partial t} = -\nabla_{\mathbf{x}} \cdot (\mathbf{v}(\mathbf{x})\rho) + \nabla\nabla : (\mathbf{D}(\mathbf{x})\rho) \tag{6}$$

Note that equation (6) is very similar to the population balance equation (PBE), except that equation (6) describes a probability density $\rho$ instead of a number-density $n$, and does not contain a source term $g$. From the theory of stochastic processes, we know that the diffusion kernel $\mathbf{D}(\mathbf{x})$ in eq. (6) is the second infinitesimal moment of the jump process defined by the reaction network in eq. (4) and can be expressed in terms of the transition rates $p_r$ as:

$$\mathbf{D}(\mathbf{x}) = \frac{1}{2}\sum_{r=1}^{R} \nu_r \nu_r^T p_r(\mathbf{x}) \tag{7}$$

Our analogue of equation (4) is the biochemical GRN described by the synthesis of mRNAs and proteins, respectively [11,12]. The ODEs describing the GRN (§1) can be decomposed into four fundamental reactions (table 3) under the assumption that the system state vector $\mathbf{x}$ can be compactly written as $\mathbf{x} = [\mathbf{mRNA}, \mathbf{protein}]^T$. The diffusion tensor can then be

Table 3: Reaction set for the mRNA-protein GRN.

| Reaction | Process | Stoichiometry ($\nu_r^T$) | Propensity ($p_r(\mathbf{x})$) |
|---|---|---|---|
| 1 | mRNA Synthesis | $[1, 0]$ | $\alpha + \gamma f(\mathbf{p})$ |
| 2 | mRNA Degradation | $[-1, 0]$ | $\delta m$ |
| 3 | Protein Translation | $[0, 1]$ | $\beta m$ |
| 4 | Protein Degradation | $[0, -1]$ | $\mu p$ |

constructed as the sum of the contributions from each reaction for each biochemical species in the GRN, thereby giving rise to a block-diagonal structure:

$$\mathbf{D}(\mathbf{x}) = \frac{1}{2}\left[\begin{array}{ccc|ccc} \alpha_1 + \gamma_1 f_1(p) + \delta m_1 & \dots & 0 & 0 & \dots & 0 \\ \vdots & \ddots & \vdots & \vdots & \ddots & \vdots \\ 0 & \dots & \alpha_{13} + \gamma_{12} f_{12}(p) + \delta m_{12} & 0 & \dots & 0 \\ \hline 0 & \dots & 0 & \beta_1 m_1 + \mu_1 p_1 & \dots & 0 \\ \vdots & \ddots & \vdots & \vdots & \ddots & \vdots \\ 0 & \dots & 0 & 0 & \dots & \beta_{13} m_{13} + \mu_{12} p_{12} \end{array}\right] \in \mathbb{R}^{24\times 24} \tag{8}$$

Note that the use of the regulatory Hill function $f(p)$ represents a quasi-steady-state approximation (QSSA) of the promoter binding kinetics. This coarse-grained approach assumes that the binding and unbinding of transcription factors occur on a significantly faster timescale than mRNA synthesis and degradation, allowing for the reduction of the promoter's discrete state transitions into a continuous, non-linear propensity function. However, we lose some of the intrinsic noise due to this coarse-graining as it doesn't take into account effects like transcriptional bursting, translational bursting or fluctuations in the number of ribosomes [13]. Furthermore, the diagonal structure of the tensor rests on the assumption that each independent reaction (transcription, translation, degradation) changes the abundance of only a single molecular species, and any correlations are absent. The inclusion of the diffusion tensor is important for maintaining a physically realistic representation of cellular heterogeneity. In its absence, the purely deterministic and contractive nature of the drift term would cause all cells to converge toward a single phenotype. Consequently, the population density would collapse into a Dirac delta distribution, failing to capture the phenotypic variance observed in biological populations [14].

### 3 Full SINDy library construction and discovered equations

#### Library construction

Given the $d$-dimensional latent state $\mathbf{z} \in \mathbb{R}^d$ and the external cytokine input $\mathbf{u} \in \mathbb{R}^m$, the SINDy framework approximates the latent-space velocity as a sparse linear combination drawn from a fixed candidate library:

$$\dot{\mathbf{z}} = \mathbf{\Theta}(\mathbf{z}, \mathbf{u})\, \mathbf{\Xi}, \qquad \mathbf{\Theta}(\mathbf{z}, \mathbf{u}) \in \mathbb{R}^{1 \times N_\theta}, \quad \mathbf{\Xi} \in \mathbb{R}^{N_\theta \times d}. \tag{9}$$

The library $\mathbf{\Theta}$ is assembled from four feature groups evaluated at each snapshot:

$$\begin{aligned}\Theta(\mathbf{z}, \mathbf{u}) = \Big\{&1,\\ &\prod_k z_{i_k} \text{ with } 1 \le |\alpha| \le p,\\ &u_j, \quad j = 0, \dots, m-1,\\ &z_i u_j, \quad i = 0, \dots, d-1,\ j = 0, \dots, m-1\Big\}.\end{aligned}$$

The full library row vector is therefore

$$\mathbf{\Theta}(\mathbf{z}, \mathbf{u}) = \Big[\underbrace{1}_{\text{const}},\ \underbrace{z_0, \dots, z_{d-1}}_{\text{deg 1}},\ \underbrace{z_0^2,\ z_0 z_1,\ \dots,\ z_{d-1}^2}_{\text{deg 2}},\ \underbrace{z_0^3,\ \dots,\ z_{d-1}^3}_{\text{deg 3}},\ \underbrace{u_0,\ \dots,\ u_{m-1}}_{\text{inputs}},\ \underbrace{z_0 u_0,\ \dots,\ z_{d-1} u_{m-1}}_{\text{cross terms}}\Big]. \tag{10}$$

The number of polynomial features of degree $k$ drawn from $d$ variables with repetition is $\binom{d+k-1}{k}$. With latent dimension $d = 5$, polynomial order $p = 3$, and input dimension $m = 2$, the total library size is

$$N_\theta = 1 + \underbrace{\binom{5}{1}}_{5} + \underbrace{\binom{6}{2}}_{15} + \underbrace{\binom{7}{3}}_{35} + m + d \cdot m = 1 + 5 + 15 + 35 + 2 + 10 = 68. \tag{11}$$

Sparsity in $\mathbf{\Xi}$ is enforced jointly through: (i) an $\ell_1$ penalty $\mathcal{L}_s = \|\mathbf{\Xi} \odot \mathbf{M}\|_1$ included in the training objective (§6), where $\mathbf{M} \in \{0,1\}^{N_\theta \times d}$ is a binary mask and (ii) a hard-thresholding step applied periodically during training that permanently zeros any entry with $|\Xi_{ij}| < \lambda$ and removes it from further gradient updates. The mask $\mathbf{M}$ is monotonically non-increasing: once an entry is zeroed it cannot recover.

#### Discovered equations

After training with the composite loss and iterative thresholding, the identified latent-space dynamics for the $d = 5$ dimensional state $\mathbf{z} = (z_0, z_1, z_2, z_3, z_4)^\top$ are as follows. The cytokine inputs $u_0$ (pro-inflammatory: IFN$\gamma$ + LPS) and $u_1$ (anti-inflammatory: IL-4 + IL-13) enter both as direct linear drivers and through state–input coupling terms.

$$\begin{aligned}\dot{z}_0 = &-0.0098 - 0.0243\, z_0 + 0.0190\, z_1 + 0.0078\, z_2 + 0.0159\, z_3 + 0.0028\, z_4\\ &- 0.0082\, z_0^2 + 0.0063\, z_0 z_1 - 0.0090\, z_0 z_2 - 0.0017\, z_0 z_3 - 0.0082\, z_0 z_4 - 0.0003\, z_1 z_2 - 0.0030\, z_1 z_3\\ &- 0.0009\, z_1 z_4 - 0.0043\, z_2^2 + 0.0044\, z_2 z_3 + 0.0036\, z_2 z_4 + 0.0067\, z_3^2 + 0.0137\, z_3 z_4 + 0.0133\, z_4^2\\ &- 0.0001\, z_0^3 - 0.0021\, z_0^2 z_1 - 0.0001\, z_0^2 z_2 + 0.0011\, z_0^2 z_3 + 0.0020\, z_0^2 z_4 + 0.0049\, z_0 z_1 z_2\\ &- 0.0007\, z_0 z_1 z_3 + 0.0018\, z_0 z_1 z_4 - 0.0019\, z_0 z_2^2 - 0.0023\, z_0 z_2 z_3 - 0.0015\, z_0 z_2 z_4\\ &- 0.0011\, z_0 z_3^2 + 0.0001\, z_0 z_3 z_4 - 0.0008\, z_0 z_4^2 + 0.0012\, z_1^3 - 0.0006\, z_1^2 z_2 - 0.0007\, z_1^2 z_4\end{aligned}$$

$$
\begin{aligned}
&- 0.0016\, z_1 z_2^2 - 0.0004\, z_1 z_2 z_3 - 0.0045\, z_1 z_2 z_4 + 0.0002\, z_1 z_3^2 + 0.0042\, z_1 z_3 z_4 - 0.0002\, z_1 z_4^2 \\
&- 0.0015\, z_2^3 - 0.0003\, z_2^2 z_4 + 0.0003\, z_2 z_3^2 - 0.0028\, z_2 z_3 z_4 - 0.0003\, z_2 z_4^2 - 0.0024\, z_3^2 z_4 - 0.0052\, z_3 z_4^2 - 0.0011\, z_4^3 \\
&- 0.0056\, u_0 + 0.0025\, u_1 + 0.0024\, z_0 u_0 - 0.0005\, z_0 u_1 - 0.0017\, z_1 u_0 + 0.0013\, z_1 u_1 \\
&- 0.0008\, z_2 u_0 - 0.0008\, z_2 u_1 + 0.0007\, z_3 u_0 + 0.0003\, z_3 u_1 - 0.0024\, z_4 u_0 \qquad (12)
\end{aligned}
$$

$$
\begin{aligned}
\dot{z}_1 = &- 0.0276 + 0.0017\, z_0 - 0.0411\, z_1 - 0.0390\, z_3 - 0.0184\, z_4 \\
&- 0.0132\, z_0^2 + 0.0050\, z_0 z_1 - 0.0086\, z_0 z_2 + 0.0008\, z_0 z_3 + 0.0060\, z_0 z_4 - 0.0005\, z_1^2 + 0.0114\, z_1 z_2 \\
&- 0.0163\, z_1 z_3 - 0.0153\, z_1 z_4 - 0.0134\, z_2^2 - 0.0004\, z_2 z_3 - 0.0060\, z_2 z_4 - 0.0062\, z_3^2 - 0.0050\, z_3 z_4 \\
&+ 0.0018\, z_0^3 - 0.0014\, z_0^2 z_1 - 0.0022\, z_0^2 z_2 - 0.0040\, z_0^2 z_3 - 0.0019\, z_0^2 z_4 - 0.0048\, z_0 z_1^2 \\
&+ 0.0004\, z_0 z_1 z_3 + 0.0052\, z_0 z_1 z_4 - 0.0006\, z_0 z_2^2 + 0.0002\, z_0 z_2 z_3 - 0.0003\, z_0 z_2 z_4 + 0.0012\, z_0 z_3 z_4 \\
&- 0.0023\, z_0 z_4^2 + 0.0011\, z_1^3 - 0.0022\, z_1^2 z_2 - 0.0039\, z_1^2 z_3 - 0.0022\, z_1^2 z_4 + 0.0024\, z_1 z_2^2 \\
&- 0.0034\, z_1 z_2 z_3 - 0.0030\, z_1 z_2 z_4 - 0.0016\, z_1 z_3 z_4 - 0.0020\, z_1 z_4^2 - 0.0026\, z_2^3 \\
&- 0.0017\, z_2^2 z_3 - 0.0030\, z_2^2 z_4 + 0.0000\, z_2 z_3 z_4 + 0.0004\, z_2 z_4^2 + 0.0019\, z_3^3 + 0.0010\, z_3^2 z_4 - 0.0011\, z_3 z_4^2 - 0.0004\, z_4^3 \\
&+ 0.0026\, u_0 - 0.0018\, z_0 u_0 + 0.0013\, z_0 u_1 - 0.0008\, z_1 u_1 + 0.0002\, z_2 u_0 - 0.0012\, z_3 u_0 - 0.0012\, z_3 u_1 + 0.0018\, z_4 u_0 + 0.0016\, z_4 u_1 \\
& \qquad (13)
\end{aligned}
$$

$$
\begin{aligned}
\dot{z}_2 = &+ 0.0240 + 0.0064\, z_0 - 0.0165\, z_1 - 0.0082\, z_2 - 0.0112\, z_3 - 0.0147\, z_4 \\
&- 0.0014\, z_0^2 - 0.0050\, z_0 z_1 + 0.0019\, z_0 z_2 + 0.0054\, z_0 z_3 + 0.0037\, z_0 z_4 - 0.0040\, z_1^2 + 0.0074\, z_1 z_2 \\
&+ 0.0041\, z_1 z_3 + 0.0047\, z_1 z_4 + 0.0050\, z_2^2 + 0.0157\, z_2 z_3 + 0.0090\, z_2 z_4 - 0.0029\, z_3^2 - 0.0072\, z_3 z_4 - 0.0067\, z_4^2 \\
&+ 0.0006\, z_0^3 + 0.0024\, z_0^2 z_1 - 0.0002\, z_0^2 z_2 - 0.0033\, z_0^2 z_3 - 0.0013\, z_0^2 z_4 - 0.0006\, z_0 z_1^2 \\
&- 0.0011\, z_0 z_1 z_2 + 0.0043\, z_0 z_1 z_3 + 0.0004\, z_0 z_1 z_4 + 0.0009\, z_0 z_2^2 - 0.0003\, z_0 z_2 z_4 - 0.0013\, z_0 z_4^2 \\
&- 0.0014\, z_1^3 - 0.0023\, z_1^2 z_2 - 0.0012\, z_1^2 z_3 + 0.0016\, z_1^2 z_4 + 0.0030\, z_1 z_2^2 + 0.0068\, z_1 z_2 z_3 + 0.0051\, z_1 z_2 z_4 \\
&- 0.0034\, z_1 z_3^2 + 0.0008\, z_1 z_3 z_4 - 0.0019\, z_1 z_4^2 - 0.0005\, z_2^2 z_3 + 0.0008\, z_2^2 z_4 + 0.0002\, z_2 z_3^2 \\
&- 0.0007\, z_2 z_3 z_4 + 0.0006\, z_2 z_4^2 + 0.0003\, z_3^3 + 0.0008\, z_3^2 z_4 + 0.0017\, z_3 z_4^2 + 0.0011\, z_4^3 \\
&+ 0.0006\, u_0 - 0.0027\, u_1 + 0.0007\, z_0 u_0 - 0.0011\, z_0 u_1 + 0.0017\, z_2 u_0 + 0.0007\, z_2 u_1 + 0.0009\, z_3 u_0 + 0.0009\, z_3 u_1 - 0.0006\, z_4 u_1 \qquad (14)
\end{aligned}
$$

$$
\begin{aligned}
\dot{z}_3 = &- 0.0226 + 0.0059\, z_0 - 0.0121\, z_1 + 0.0293\, z_2 + 0.0132\, z_3 - 0.0119\, z_4 \\
&+ 0.0007\, z_0^2 + 0.0040\, z_0 z_1 + 0.0109\, z_0 z_2 - 0.0083\, z_0 z_3 + 0.0054\, z_0 z_4 - 0.0096\, z_1^2 - 0.0121\, z_1 z_2 \\
&- 0.0011\, z_1 z_3 + 0.0039\, z_1 z_4 - 0.0056\, z_2^2 + 0.0071\, z_2 z_3 + 0.0033\, z_2 z_4 + 0.0006\, z_3^2 + 0.0056\, z_3 z_4 - 0.0117\, z_4^2 \\
&+ 0.0010\, z_0^2 z_1 + 0.0010\, z_0^2 z_2 - 0.0008\, z_0^2 z_3 + 0.0012\, z_0^2 z_4 + 0.0019\, z_0 z_1^2 + 0.0002\, z_0 z_1 z_2 \\
&- 0.0019\, z_0 z_1 z_3 - 0.0008\, z_0 z_1 z_4 + 0.0009\, z_0 z_2^2 + 0.0010\, z_0 z_2 z_3 - 0.0025\, z_0 z_2 z_4 - 0.0012\, z_0 z_3^2 \\
&+ 0.0024\, z_0 z_3 z_4 - 0.0026\, z_0 z_4^2 - 0.0003\, z_1^3 - 0.0011\, z_1^2 z_2 + 0.0010\, z_1^2 z_4 - 0.0002\, z_1 z_2^2 + 0.0030\, z_1 z_2 z_3 \\
&+ 0.0043\, z_1 z_2 z_4 - 0.0032\, z_1 z_3^2 + 0.0001\, z_1 z_3 z_4 + 0.0001\, z_1 z_4^2 - 0.0010\, z_2^3 + 0.0009\, z_2^2 z_3 \\
&- 0.0002\, z_2 z_3^2 + 0.0015\, z_2 z_3 z_4 - 0.0012\, z_2 z_4^2 - 0.0014\, z_3^3 + 0.0002\, z_3^2 z_4 - 0.0003\, z_3 z_4^2 + 0.0014\, z_4^3 \\
&+ 0.0009\, u_0 - 0.0032\, u_1 - 0.0005\, z_0 u_0 - 0.0001\, z_0 u_1 + 0.0013\, z_1 u_0 - 0.0035\, z_1 u_1 + 0.0011\, z_2 u_0 - 0.0025\, z_2 u_1 \\
&+ 0.0009\, z_3 u_1 + 0.0019\, z_4 u_0 - 0.0013\, z_4 u_1 \qquad (15)
\end{aligned}
$$

$$
\begin{aligned}
\dot{z}_4 = &- 0.0394 - 0.0170\, z_0 - 0.0109\, z_1 + 0.0165\, z_2 - 0.0159\, z_3 - 0.0193\, z_4 \\
&- 0.0065\, z_0^2 + 0.0144\, z_0 z_1 - 0.0022\, z_0 z_3 - 0.0028\, z_0 z_4 - 0.0090\, z_1^2 + 0.0001\, z_1 z_2 - 0.0128\, z_1 z_3 - 0.0062\, z_1 z_4 \\
&- 0.0029\, z_2^2 + 0.0006\, z_2 z_3 + 0.0012\, z_2 z_4 + 0.0014\, z_3^2 - 0.0002\, z_3 z_4 \\
&+ 0.0009\, z_0^3 - 0.0016\, z_0^2 z_1 - 0.0001\, z_0^2 z_2 - 0.0012\, z_0^2 z_3 - 0.0004\, z_0 z_1^2 + 0.0003\, z_0 z_1 z_2 \\
&- 0.0006\, z_0 z_1 z_3 + 0.0006\, z_0 z_1 z_4 + 0.0019\, z_0 z_2^2 - 0.0031\, z_0 z_2 z_3 - 0.0002\, z_0 z_2 z_4 - 0.0023\, z_0 z_3 z_4 - 0.0003\, z_0 z_4^2 \\
&- 0.0031\, z_1^2 z_2 - 0.0009\, z_1^2 z_3 - 0.0009\, z_1^2 z_4 + 0.0002\, z_1 z_2^2 - 0.0033\, z_1 z_2 z_3 + 0.0006\, z_1 z_2 z_4 \\
&- 0.0015\, z_1 z_3 z_4 + 0.0002\, z_1 z_4^2 + 0.0005\, z_2^2 z_3 + 0.0010\, z_2 z_3^2 + 0.0015\, z_2 z_3 z_4 - 0.0008\, z_2 z_4^2 \\
&+ 0.0008\, z_3^3 - 0.0015\, z_3^2 z_4 - 0.0009\, z_4^3 \\
&+ 0.0006\, u_0 - 0.0016\, z_0 u_0 - 0.0006\, z_1 u_0 + 0.0007\, z_1 u_1 - 0.0019\, z_2 u_0 + 0.0006\, z_2 u_1 \\
&- 0.0013\, z_3 u_0 - 0.0003\, z_3 u_1 + 0.0001\, z_4 u_0 + 0.0015\, z_4 u_1 \qquad (16)
\end{aligned}
$$

### 4 PBE transformation under autoencoder

Let the original state variable be $\mathbf{x}_t \in \mathbb{R}^N$. Suppose the particle dynamics in the original state space are governed by the Itô SDE:

$$d\mathbf{x}_t = \mathbf{v}(\mathbf{x}_t, t)\, dt + \mathbf{B}(\mathbf{x}_t, t)\, d\mathbf{W}_t,$$

where $\mathbf{v}(\mathbf{x}, t) \in \mathbb{R}^N$ is the drift vector, $\mathbf{B}(\mathbf{x}, t) \in \mathbb{R}^{N\times r}$ is the noise-amplitude matrix, and $\mathbf{W}_t \in \mathbb{R}^r$ is a standard Wiener process. The corresponding diffusion tensor in the original state space is

$$\mathbf{D}(\mathbf{x}, t) = \mathbf{B}(\mathbf{x}, t)\mathbf{B}^T(\mathbf{x}, t).$$

Now define the latent variable $\mathbf{z}_t \in \mathbb{R}^d$ through an encoder map $\mathbf{z}_t = \mathcal{E}(\mathbf{x}_t)$, where $\mathcal{E} : \mathbb{R}^N \to \mathbb{R}^d$. Writing the encoder componentwise,

$$\mathcal{E}(\mathbf{x}) = \begin{bmatrix} \mathcal{E}_1(\mathbf{x}) \\ \mathcal{E}_2(\mathbf{x}) \\ \vdots \\ \mathcal{E}_d(\mathbf{x}) \end{bmatrix},$$

we have $z_{k,t} = \mathcal{E}_k(\mathbf{x}_t)$, $k = 1, \ldots, d$. By **Itô's lemma** [15], for each component $\mathcal{E}_k(\mathbf{x}_t)$,

$$dz_{k,t} = \nabla\mathcal{E}_k(\mathbf{x}_t)^T d\mathbf{x}_t + \frac{1}{2}\,\mathrm{Tr}\left[\mathbf{D}(\mathbf{x}_t, t)\nabla^2\mathcal{E}_k(\mathbf{x}_t)\right] dt.$$

Substituting the original SDE gives,

$$dz_{k,t} = \nabla\mathcal{E}_k(\mathbf{x}_t)^T \left[\mathbf{v}(\mathbf{x}_t, t)\, dt + \mathbf{B}(\mathbf{x}_t, t)\, d\mathbf{W}_t\right] + \frac{1}{2}\,\mathrm{Tr}\left[\mathbf{D}(\mathbf{x}_t, t)\nabla^2\mathcal{E}_k(\mathbf{x}_t)\right] dt.$$

Therefore,

$$dz_{k,t} = \left[\nabla\mathcal{E}_k(\mathbf{x}_t)^T\mathbf{v}(\mathbf{x}_t, t) + \frac{1}{2}\,\mathrm{Tr}\left[\mathbf{D}(\mathbf{x}_t, t)\nabla^2\mathcal{E}_k(\mathbf{x}_t)\right]\right] dt + \nabla\mathcal{E}_k(\mathbf{x}_t)^T\mathbf{B}(\mathbf{x}_t, t)\, d\mathbf{W}_t.$$

Thus, the transformed latent-space SDE can be written as

$$d\mathbf{z}_t = \tilde{\mathbf{v}}(\mathbf{z}_t, t)\, dt + \tilde{\mathbf{B}}(\mathbf{z}_t, t)\, d\mathbf{W}_t.$$

The encoder Jacobian is

$$\mathbf{J}_{\mathcal{E}}(\mathbf{x}) = \frac{\partial\mathcal{E}}{\partial\mathbf{x}} \in \mathbb{R}^{d\times N}.$$

The latent-space diffusion tensor is therefore

$$\tilde{\mathbf{D}}(\mathbf{z}, t) = \left[\mathbf{J}_{\mathcal{E}}(\mathbf{x})\mathbf{D}(\mathbf{x}, t)\mathbf{J}_{\mathcal{E}}^T(\mathbf{x})\right]_{\mathbf{x}=\mathcal{D}(\mathbf{z})}.$$

And the latent-space drift vector is

$$\tilde{\mathbf{v}}(\mathbf{z}, t) = \left[\mathbf{J}_{\mathcal{E}}(\mathbf{x})\mathbf{v}(\mathbf{x}, t) + \frac{1}{2}\boldsymbol{c}_{\mathcal{E}}(\mathbf{x}, t)\right]_{\mathbf{x}=\mathcal{D}(\mathbf{z})},$$

where the Itô correction vector $\boldsymbol{c}_{\mathcal{E}}(\mathbf{x}, t) \in \mathbb{R}^d$ is defined componentwise as

$$\left[\boldsymbol{c}_{\mathcal{E}}(\mathbf{x}, t)\right]_k = \mathrm{Tr}\left[\mathbf{D}(\mathbf{x}, t)\nabla^2\mathcal{E}_k(\mathbf{x})\right], \qquad k = 1, \ldots, d.$$

Therefore, the transformed SDE is

$$\boxed{d\mathbf{z}_t = \left[\mathbf{J}_{\mathcal{E}}(\mathbf{x}_t)\mathbf{v}(\mathbf{x}_t, t) + \frac{1}{2}\mathbf{c}_{\mathcal{E}}(\mathbf{x}_t, t)\right] dt + \mathbf{J}_{\mathcal{E}}(\mathbf{x}_t)\mathbf{B}(\mathbf{x}_t, t)d\mathbf{W}_t.}$$

From the latent-space SDE, $d\mathbf{z}_t = \tilde{\mathbf{v}}(\mathbf{z}_t, t)\, dt + \tilde{\mathbf{B}}(\mathbf{z}_t, t)\, d\mathbf{W}_t$, the corresponding Kramers–Moyal coefficients are $\mathbf{a}^{(1)}(\mathbf{z}, t) = \tilde{\mathbf{v}}(\mathbf{z}, t)$ and $\mathbf{a}^{(2)}(\mathbf{z}, t) = \tilde{\mathbf{D}}(\mathbf{z}, t) = \tilde{\mathbf{B}}(\mathbf{z}, t)\tilde{\mathbf{B}}^T(\mathbf{z}, t)$. Therefore, the latent-space number density $\tilde{n}(\mathbf{z}, t)$ satisfies the Fokker–Planck type, or population balance, equation

$$\frac{\partial\tilde{n}(\mathbf{z}, t)}{\partial t} = -\sum_{i=1}^{n_z}\frac{\partial}{\partial z_i}\left[\tilde{v}_i(\mathbf{z}, t)\tilde{n}(\mathbf{z}, t)\right] + \frac{1}{2}\sum_{i=1}^{n_z}\sum_{j=1}^{n_z}\frac{\partial^2}{\partial z_i\partial z_j}\left[\tilde{D}_{ij}(\mathbf{z}, t)\tilde{n}(\mathbf{z}, t)\right].$$

If a source or sink term is included, the latent-space PBE becomes

$$\frac{\partial \tilde{n}(\mathbf{z},t)}{\partial t} = -\sum_{i=1}^{n_z} \frac{\partial}{\partial z_i}\left[\tilde{v}_i(\mathbf{z},t)\tilde{n}(\mathbf{z},t)\right] + \frac{1}{2}\sum_{i=1}^{n_z}\sum_{j=1}^{n_z} \frac{\partial^2}{\partial z_i \partial z_j}\left[\tilde{D}_{ij}(\mathbf{z},t)\tilde{n}(\mathbf{z},t)\right] + \tilde{S}(\tilde{n},\mathbf{z},t).$$

Or in compact vector notation,

$$\boxed{\frac{\partial \tilde{n}}{\partial t} = -\nabla_{\mathbf{z}} \cdot \left[\left(\mathbf{J}_{\mathcal{E}}\mathbf{v} + \frac{1}{2}\boldsymbol{c}_{\mathcal{E}}\right)\tilde{n}\right] + \frac{1}{2}\nabla_{\mathbf{z}}\nabla_{\mathbf{z}} : \left[\left(\mathbf{J}_{\mathcal{E}}\mathbf{D}\mathbf{J}_{\mathcal{E}}^{T}\right)\tilde{n}\right] + \tilde{S}.}$$

Where, $[\boldsymbol{c}_{\mathcal{E}}]_k = \mathrm{Tr}\left[\mathbf{D}\nabla^2\mathcal{E}_k\right]$.

### 5 Latent-space stochastic particle update

The latent-space PBE is to be solved numerically within the latent space of the autoencoder (AE). Because the AE is nonlinear, the geometry of the learned latent manifold is *a priori* unknown and may be highly curved. This poses a significant challenge for traditional grid-based approaches, such as finite volume or finite difference methods, which require a well-defined mesh and explicit boundary conditions. Moreover, the latent coordinates are non-physical variables and therefore do not admit straightforward biological boundary conditions. These computational and geometric constraints motivate the use of a mesh-free particle method. We note that the conservative drift–diffusion part of the latent-space PBE admits the following microscopic stochastic representation:

$$d\mathbf{Z}_t = \tilde{\mathbf{v}}(\mathbf{Z}_t,t)\,dt + \tilde{\mathbf{B}}(\mathbf{Z}_t,t)\,d\mathbf{W}_t, \tag{17}$$

where $\mathbf{Z}_t \in \mathbb{R}^d$ is a stochastic particle trajectory in the $d$-dimensional latent space, $d\mathbf{W}_t$ is a standard vector Wiener process, and the noise-amplitude matrix $\tilde{\mathbf{B}}$ satisfies

$$\frac{1}{2}\tilde{\mathbf{B}}(\mathbf{z},t)\tilde{\mathbf{B}}(\mathbf{z},t)^T = \tilde{\mathbf{D}}(\mathbf{z},t). \tag{18}$$

Thus, rather than discretizing the latent number density ($\tilde{n}$) on a mesh, we can approximate it by an ensemble of particles,

$$\tilde{n}(\mathbf{z},t) \approx \sum_{p=1}^{N_p} w_p(t)\,\delta_D\left(\mathbf{z} - \mathbf{Z}_t^{(p)}\right), \tag{19}$$

where $\mathbf{Z}_t^{(p)}$ is the position of particle $p$, $w_p(t)$ is its weight, and $N_p$ is the total number of particles [16]. In the baseline case considered here, we set $S_z(n_z,\mathbf{z},t) = 0$, so that no local source, death, or growth term is included. Consequently, the particle weights remain constant:

$$w_p(t) = w_p(0), \qquad p = 1,\ldots,N_p. \tag{20}$$

Under this assumption, the stochastic particle ensemble evolves only through drift and diffusion in latent space and the particle trajectories are advanced using the Euler–Maruyama discretization of (17) as:

$$\mathbf{Z}_{n+1}^{(p)} = \mathbf{Z}_n^{(p)} + \Delta t\,\tilde{\mathbf{v}}\left(\mathbf{Z}_n^{(p)},t_n\right) + \sqrt{\Delta t}\,\tilde{\mathbf{B}}\left(\mathbf{Z}_n^{(p)},t_n\right)\boldsymbol{\xi}_n^{(p)}, \tag{21}$$

where $\boldsymbol{\xi}_n^{(p)} \sim \mathcal{N}(\mathbf{0},\mathbf{I}_d)$ is a standard Gaussian random vector in the latent space. An important distinction between the physical-space PBE and the latent-space PBE is that the latent-space transport coefficients are not available in closed form. In the original physical state space, the drift and diffusion are prescribed by the mechanistic gene-regulatory model. However, after transformation through the nonlinear encoder $\mathbf{z} = \mathcal{E}(\mathbf{x})$, the corresponding latent-space PBE coefficients depend on derivatives of the trained autoencoder. Therefore, the coefficients in the latent PBE, or equivalently in the latent SDE, are evaluated numerically at each particle location at each time-step as per Algorithm-1.

### 6 Training losses

**Reconstruction loss $\mathcal{L}_r$.** The encoder compresses a 24-dimensional gene expression state into $d$ numbers; the decoder attempts to recover the original 24 dimensions from those $d$ numbers plus the cytokine input. $\mathcal{L}_r$ is the mean squared error of this round-trip, penalizing any information destroyed during compression. Minimizing it forces the latent coordinates to retain all gene-expression variation present in the data.

**Algorithm 1** Latent-space stochastic particle update with Itô correction

**input:** particles $\{\mathbf{Z}_n^{(p)}\}_{p=1}^{N_p}$, encoder $\mathcal{E}$, decoder $\mathcal{D}$, physical drift $\mathbf{v}_x$, physical diffusion $\mathbf{D}_x$, timestep $\Delta t$
**for** $p = 1$ to $N_p$ **do**
  $\mathbf{X}_n^{(p)} = \mathcal{D}(\mathbf{Z}_n^{(p)})$ ▷ decode particle
  $\mathbf{v}_x^{(p)} = \mathbf{v}_x\left(\mathbf{X}_n^{(p)}; u_{\text{pro}}, u_{\text{anti}}\right)$ ▷ evaluate physical drift
  $\mathbf{D}_x^{(p)} = \mathbf{D}_x\left(\mathbf{X}_n^{(p)}\right)$ ▷ evaluate physical diffusion
  $\mathbf{J}_{\mathcal{E}}^{(p)} = \left.\frac{\partial \mathcal{E}}{\partial \mathbf{x}}\right|_{\mathbf{x}=\mathbf{X}_n^{(p)}}$ ▷ evaluate encoder Jacobian
  $H_{\mathcal{E}_i}^{(p)} = \left.\frac{\partial^2 \mathcal{E}_i}{\partial \mathbf{x}^2}\right|_{\mathbf{x}=\mathbf{X}_n^{(p)}}, \quad i = 1, \ldots, d$ ▷ evaluate encoder Hessians
  $\tilde{\mathbf{v}}_z^{(p)} = \mathbf{J}_{\mathcal{E}}^{(p)} \mathbf{v}_x^{(p)}$ ▷ push forward physical drift
  $c_{\mathcal{E},i}^{(p)} = \mathrm{Tr}\left[\mathbf{D}_x^{(p)} H_{\mathcal{E}_i}^{(p)}\right], \quad i = 1, \ldots, d$ ▷ compute Itô correction
  $\mathbf{c}_{\mathcal{E}}^{(p)} = \left[c_{\mathcal{E},1}^{(p)}, \ldots, c_{\mathcal{E},k}^{(p)}\right]^T$ ▷ assemble correction vector
  $\tilde{\mathbf{v}}^{(p)} = \tilde{\mathbf{v}}_z^{(p)} + \mathbf{c}_{\mathcal{E}}^{(p)}$ ▷ assemble complete latent drift
  $\tilde{\mathbf{D}}^{(p)} = \mathbf{J}_{\mathcal{E}}^{(p)} \mathbf{D}_x^{(p)} \left(\mathbf{J}_{\mathcal{E}}^{(p)}\right)^T$ ▷ push forward diffusion
  find $\tilde{\mathbf{B}}^{(p)}$ such that

$$\tilde{\mathbf{B}}^{(p)} \left(\tilde{\mathbf{B}}^{(p)}\right)^T = 2\tilde{\mathbf{D}}^{(p)}$$

▷ noise amplitude
  $\boldsymbol{\xi}_n^{(p)} \sim \mathcal{N}(\mathbf{0}, \mathbf{I}_d)$ ▷ sample latent noise
  $\mathbf{Z}_{n+1}^{(p)} = \mathbf{Z}_n^{(p)} + \Delta t\, \tilde{\mathbf{v}}^{(p)} + \sqrt{\Delta t}\, \tilde{\mathbf{B}}^{(p)} \boldsymbol{\xi}_n^{(p)}$ ▷ Euler–Maruyama update
**end for**
**output:** updated particles $\{\mathbf{Z}_{n+1}^{(p)}\}_{p=1}^{N_p}$

**Dynamics consistency loss $\mathcal{L}_d$.** At every snapshot we have the observed rate of change $\dot{\mathbf{x}}$ from the GRN ODE. Pushing this through the encoder Jacobian gives the "true" latent velocity $J_E\, \dot{\mathbf{x}}$. $\mathcal{L}_d$ is the squared error between this true latent velocity and the velocity predicted by the SINDy model $\Theta\Xi$. It enforces that the sparse equations discovered in latent space are actually consistent with how the system moves.

**Derivative reconstruction loss $\mathcal{L}_\partial$.** The inverse check of $\mathcal{L}_d$: take the SINDy-predicted latent velocity, push it back through the decoder Jacobian, and compare with the original $\dot{\mathbf{x}}$. Together $\mathcal{L}_d$ and $\mathcal{L}_\partial$ form a closed cycle as encoder and decoder must agree on velocities in both directions simultaneously.

**One-step prediction loss $\mathcal{L}_n$.** A single explicit Euler step from $\mathbf{z}$ using the predicted velocity, decoded back to state space, should land at the observed next snapshot $\mathbf{x}_{t+\Delta t}$. This loss provides a direct temporal signal that $\mathcal{L}_d$ and $\mathcal{L}_\partial$ alone cannot supply, since those terms constrain instantaneous velocities but not finite-time trajectories.

**Sparsity loss $\mathcal{L}_s$.** An $\ell_1$ penalty on the active entries of $\Xi$. Small coefficients are continuously pushed toward zero throughout training, complementing the hard thresholding step applied every 50 epochs. The combined effect is that only terms with a sufficiently large and persistent contribution survive, yielding interpretable closed-form equations.

**Fixed-point pinning loss $\mathcal{L}_{\text{fp}}$.** A stable steady state of the GRN is a gene expression pattern that does not change over time. When encoded, it should also be stationary in latent space, i.e. the SINDy velocity field should evaluate to zero there. $\mathcal{L}_{\text{fp}}$ directly penalizes any nonzero latent velocity at the encoded positions of the precomputed GRN attractors, anchoring the learned dynamics to the known biology.

**Phenotype separation loss $\mathcal{L}_{\text{sep}}$.** Under the same cytokine input the GRN can settle into either an M1 (pro-inflammatory) or M2 (anti-inflammatory) steady state — two physically distinct attractors. Nothing in $\mathcal{L}_r$ prevents the encoder from mapping both to the same latent point, which would make bistability invisible. $\mathcal{L}_{\text{sep}}$ imposes a minimum distance $m$ between every M1–M2 pair of encoded steady states, forcing the two attractors into geometrically distinct latent regions so the learned coordinate system reflects the multistable topology of the GRN.
